# A comprehensive mechanosensory connectome reveals a somatotopically organized neural circuit architecture controlling stimulus-aimed grooming of the *Drosophila* head

**DOI:** 10.1101/2025.05.19.654894

**Authors:** Steven A. Calle-Schuler, Alexis E. Santana-Cruz, Lucia Kmecová, Stefanie Hampel, Andrew M. Seeds

## Abstract

Animals respond to tactile stimulations of the body with location-appropriate behavior, such as aimed grooming. These responses are mediated by mechanosensory neurons distributed across the body, whose axons project into somatotopically organized brain regions corresponding to body location. How mechanosensory neurons interface with brain circuits to transform mechanical stimulations into location-appropriate behavior is unclear. We previously described the somatotopic organization of bristle mechanosensory neurons (BMNs) around the *Drosophila* head that elicit a sequence of location-aimed grooming movements (Eichler et al., 2024). Here, we use a serial section electron microscopy reconstruction of a full adult fly brain to identify nearly all of BMN pre- and postsynaptic partners, uncovering circuit pathways that control head grooming. Postsynaptic partners dominate the connectome, and are both excitatory and inhibitory. We identified an excitatory cholinergic hemilineage (hemilineage 23b), a developmentally related group of neurons that elicits aimed head grooming and exhibit differential connectivity with BMNs from distinct head locations, revealing a lineage-based somatotopically organized parallel circuit architecture. Presynaptic partners provide extensive BMN presynaptic inhibition, consistent with models of sensory gain control as a mechanism of suppressing grooming movements and controlling the sequence. This work provides the first comprehensive map of a somatotopically organized connectome, and reveals how this organization could shape grooming. It also reveals the mechanosensory interface with the brain, illuminating fundamental features of mechanosensory processing, including feedforward excitation and inhibition, feedback inhibition, somatotopic circuit organization, and developmental origins.

## Introduction

The motivation for this work came from our studies of the neural circuit mechanisms that underlie complex sequential behavior. The capacity to perform complex behaviors by assembling different movements in sequence is crucial for adaptive behavior and ensuring survival. A prominent conceptual framework known as the “parallel model” elucidates the neural mechanisms underlying movement sequence generation. This model posits that premotor elements corresponding to individual movements scheduled for sequential execution are simultaneously activated, followed by a sequential selection process (Bohland et al., 2010; Bullock, 2004; Houghton and Hartley, 1995; Lashley, 1951). Central to the model is a parallel circuit architecture, wherein all alternative movements intended for sequential selection are activated simultaneously and compete for output. Performance order is established through suppression, wherein earlier movements inhibit later ones within the circuit architecture. Empirical support for this model is derived from physiological and behavioral evidence spanning diverse animal species, lending credence to its universality and applicability (Averbeck et al., 2002; Mushiake et al., 2006; Seeds et al., 2014). Nonetheless, a comprehensive elucidation of the underlying neural circuits remains an ongoing pursuit.

Investigations into the grooming behavior of fruit flies (*Drosophila melanogaster*) offer insights into the circuit mechanisms underlying movement sequences. Coating flies with dust elicits a cleaning sequence that commences with the grooming of various head locations such as the eyes, proboscis, and antennae, before progressing to body locations including the abdomen, wings, and thorax (Mueller et al., 2019; Phillis et al., 1993; Seeds et al., 2014). This sequence is orchestrated by a mechanism consistent with a parallel model (Seeds et al., 2014). Essentially, different mutually exclusive grooming movements, aimed at specific head or body locations, are simultaneously activated by the presence of dust. The ensuing competition among these movements is resolved through hierarchical suppression. For instance, grooming of the eyes takes precedence, since it suppresses grooming elsewhere on the head and body. This parallel model of hierarchical suppression provides a conceptual framework for elucidating the neural circuitry governing *Drosophila* grooming (**Figure 1 – figure supplement 1A,B**).

Previous studies focused on the first layer of the grooming circuit architecture: mechanosensory neurons. Mechanosensory structures are dispersed across the surface of the head and body that respond to mechanical stimuli and elicit grooming. Among these structures, bristles are the most prevalent. Displacement of single bristles elicits grooming, during which the legs are aimed towards the stimulated area (Corfas and Dudai, 1989; Page and Matheson, 2004; Vandervorst and Ghysen, 1980). A single bristle mechanosensory neuron (BMN) innervates each bristle, becoming activated in response to the displacement of the corresponding bristle (Corfas and Dudai, 1990; Tuthill and Wilson, 2016a; Walker et al., 2000). As a result, specific aimed movements can be associated with bristles and their corresponding BMNs. Other mechanosensory structures, such as stretch receptors and chordotonal organs, also elicit grooming responses aimed at the location of the stimulus (Hampel et al., 2015, 2017, 2020a; Zhang et al., 2020).

Optogenetic activation of mechanosensory neurons simultaneously across the body elicits sequential grooming that mirrors the order of the natural sequence induced by dust (Hampel et al., 2017; Zhang et al., 2020). Hence, parallel mechanosensory pathways induce this sequence, each eliciting a movement to groom a specific location on the head or body (**Figure 1 – figure supplement 1A,B**).

The parallel model predicts different circuit features underlying the grooming sequence (Seeds et al., 2014). One is a somatotopic mechanosensory circuit architecture that elicits aimed grooming of specific locations. Indeed, the BMN axon projections in the central nervous system (CNS) show a somatotopic arrangement, where distinct projection zones—spatially localized regions of axonal arborization and synaptic output—correspond to specific head and body locations (Eichler et al., 2024; Johnson and Murphey, 1985; Murphey et al., 1989; Newland, 1991; Newland et al., 2000; Tsubouchi et al., 2017). These parallel-projecting mechanosensory neurons are hypothesized to connect with circuits that elicit grooming of those locations (**Figure 1 – figure supplement 1A-C**). In support of this, mechanosensory-connected neural circuitry has been identified that elicits aimed grooming of specific head and body locations (Hampel et al., 2020b, 2015; Yoshikawa et al., 2024; Zhang and Simpson, 2022). The model also features a hierarchical suppression mechanism among all mutually exclusive movements to be performed in the sequence, where earlier movements inhibit later ones. Activity gradients among the parallel circuits determine movement order, possibly regulated by presynaptic sensory gain control (Seeds et al., 2014).

We focus on mechanosensory pathways that elicit grooming of different locations on the *Drosophila* head. Dust-induced head grooming is performed by the forelegs that start with the eyes and progress to other locations such as the proboscis and antennae (major head locations shown in **Figure 1 – figure supplement 1C**) (Seeds et al., 2014). The primary mechanosensory structures on the head that could detect the dust are populations of bristles on the eyes, antennae, proboscis, and other head locations (**Figure 1A,B**, **Figure 1 – figure supplement 1D-G**). Each population is innervated by specific BMN types that elicit aimed grooming of their corresponding bristle locations (Eichler et al., 2024; Hampel et al., 2017; Zhang et al., 2020). We previously identified and reconstructed nearly all BMNs from around the head in a serial section electron microscopy (EM) volume of a full adult fly brain (FAFB) and mapped their distinct projections into the CNS (Eichler et al., 2024). The BMNs project in a somatotopic arrangement, wherein types innervating neighboring bristles project to overlapping zones, while those innervating distant bristles project to distinct zones (**Figure 1A-D**, **Figure 1 – figure supplement 2A-E**). Preliminary, connectomic analysis revealed that neighboring BMNs show higher postsynaptic connectivity similarity than distant BMNs (Eichler et al., 2024), consistent with the hypothesized parallel postsynaptic circuit architecture underlying grooming.

**Figure 1.**
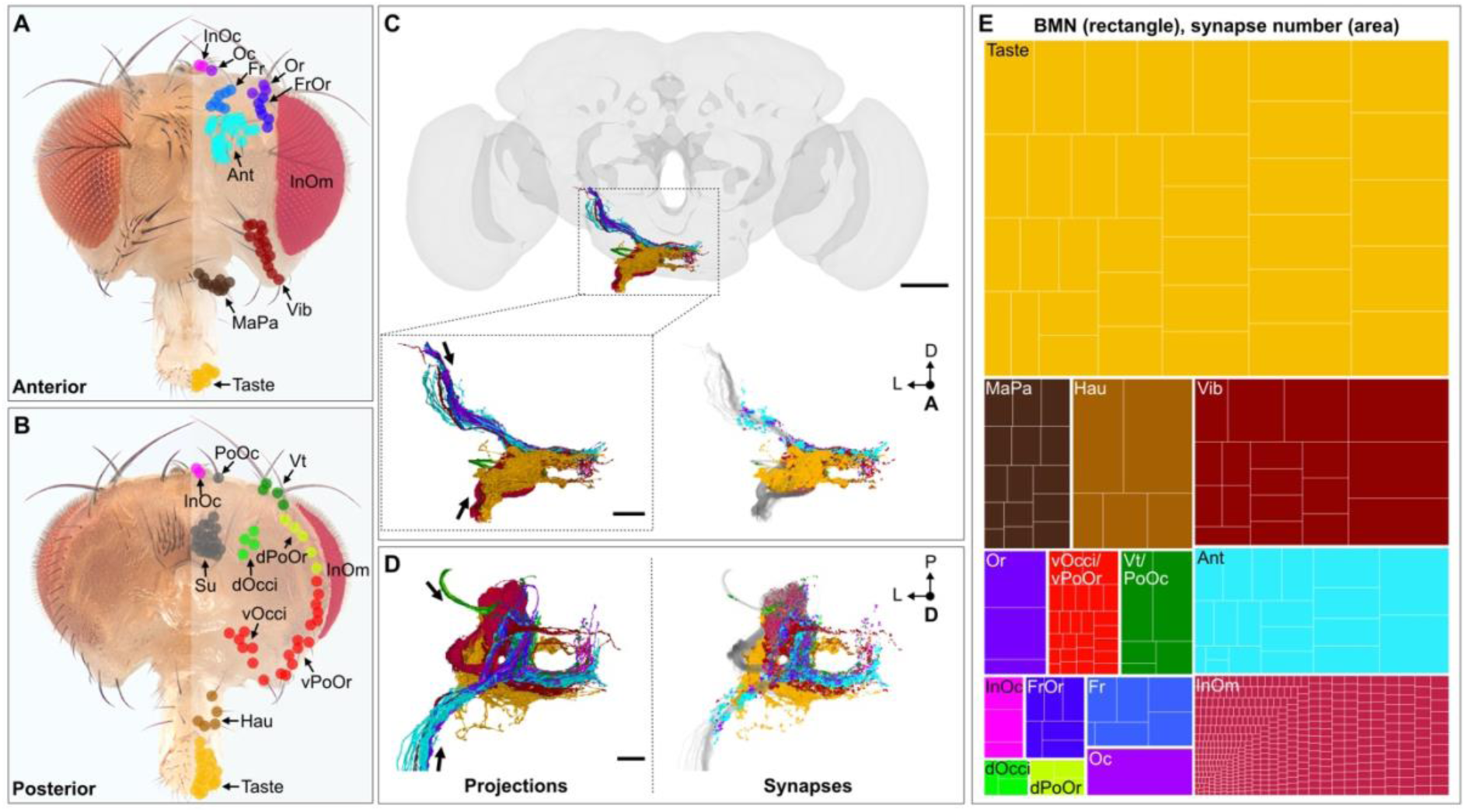
*Drosophila melanogaster* head bristle mechanosensory neuron (BMN) projections and their synapses. (**A,B**) Bristles of the anterior (**A**) and posterior (**B**) head. Color-coded dots on the right indicate bristle classifications whose names are abbreviated (full names below). Dorsal and ventral views are shown in Figure 1 **– figure supplement 1**. Su bristles are classified, but no associated BMNs have been identified. (**C,D**) Reconstructed BMN projections in the ventral brain (left, previously described in (Eichler et al., 2024)) and their corresponding pre- and postsynaptic sites (right, this study), colored by type according to the bristles that they innervate. Shown are the anterior (**C**) and dorsal (**D**) views. Overlapping projection zones are evident where synapses of different BMN types spatially intermingle, whereas segregated zones show little or no color mixing. Arrows indicate the projection directions for each incoming BMN nerve bundle. Note: BMNs are from the anatomical left side of the head, but are displayed inverted on the right as previously described (Schlegel et al., 2024). Scale bars, 50 μm (full brain) and 20 μM (anterior and dorsal zoom views). Medial and lateral views of the projections and synapses are shown in Figure 1 **– figure supplement 2** and Figure 1 **– figure supplement 3**, respectively. (**E**) Relative numbers of total synapses for each head BMN type. BMNs are named according to the bristle population that they innervate (e.g. BM-Taste neurons innervate Taste bristles). Rectangles correspond to individual BMNs whose relative areas indicate the number of total pre- and postsynaptic sites. Colors indicate individual BMNs of the same type. Underlying data is in **Supplementary file 1**. Abbreviations used to identify the bristles (**A**) and BMNs (**E**) are as follows: antennal (Ant), frontal (Fr), orbital (Or), frontoorbital (FrOr), ocellar (Oc), interocellar (InOc), vibrissae (Vib), vertical (Vt), dorsal occipital (dOcci), dorsal postorbital (dPoOr), interommatidial (InOm), ventral occipital (vOcci), ventral postorbital (vPoOr), Taste (Taste), haustellum (Hau), and maxillary palp (MaPa). Panels **A** and **B** were reproduced under the terms of the CCBY license from Figure 1A,B of Eichler *et al*. (Eichler et al., 2024).

Here, we define the synaptic connectivity of head BMNs by mapping nearly all of their pre- and postsynaptic partners—including other BMNs, ascending and descending neurons, interneurons, and motor neurons—within the FAFB dataset. Consistent with a parallel model, we find that both presynaptic and postsynaptic partners are somatotopically organized, preserving the spatial layout of the bristle map and revealing a set of parallel mechanosensory pathways that correspond to distinct head regions. Within the postsynaptic population, we identify the developmentally-related cholinergic hemilineage 23b (LB23), whose members exhibit region-specific BMN connectivity and include neurons previously shown to elicit aimed head grooming movements when activated. This demonstrates how LB23 neurons participate in parallel postsynaptic pathways that may drive discrete components of head grooming. On the input side, BMNs receive substantial presynaptic inhibition from predominantly GABAergic partners, providing strong feedback and feedforward control over mechanosensory signaling. This inhibitory architecture is consistent with hierarchical-suppression models in which inhibition regulates sensory gain and prioritizes competing actions in the grooming sequence. Together, this mechanosensory connectome reveals core organizational principles—parallel somatotopic architecture, region-specific excitatory pathways, and strong inhibitory regulation—that are thought to constitute foundational circuit motifs supporting head grooming.

## Results

### BMN synapses are somatotopically distributed in the ventral brain

In prior work (Eichler et al., 2024), we showed that head bristle populations are innervated by specific BMN types whose axons project to distinct, spatially localized regions (projection zones) in the ventral brain (**Figure 1C,D**, left, **Figure 1 – figure supplement 2A-E**). This was determined using dye fills and light-microscopy-based tracing to identify BMN types innervating defined head bristle populations and to establish their characteristic brain projection morphologies. Bristle population counts and their variability across individuals provided expectations for BMN number per type. This quantitative constraint, combined with the highly stereotyped projection morphologies, provided a correlative anatomical framework to locate and reconstruct nearly all BMNs in the FAFB serial-section EM volume and map their projections into the CNS. Because FAFB does not include the head cuticular bristles, individual BMNs could not be linked to single bristles. Therefore, these assignments are necessarily correlative and provide type-level (population) rather than single-bristle resolution. Nevertheless, this level of resolution was sufficient to define somatotopically organized projection zones.

These projection zones are also apparent at the synaptic level by comparing the spatial distributions of all BMN synapses identified in the FAFB dataset (Buhmann et al., 2021; Dorkenwald et al., 2022). Synapses of different BMN types exhibited distinct spatial distributions along their axonal projections. Segregation between projection zones is apparent where synapses of distinct BMN types occupy non-overlapping regions with little or no color mixing, whereas overlap between projection zones is visible as spatial intermixing of differently colored synapses from neighboring BMN types (**Figure 1C,D**, right, **Figure 1 – figure supplement 3A-E**).

### BMN synapses show large quantitative variation across types

Each BMN type contributed a distinct number of synapses, reflecting differences in BMN numbers and their average synapse counts (**Figure 1E**, **Supplementary file 1**). Here and throughout the study, synapse counts are based on FlyWire/Codex annotations and report individual synaptic contacts (incoming or outgoing connections), not presynaptic active sites (T-bars); thus, presynaptic counts reflect polyadic connectivity (Schlegel et al., 2024). Some BMN types varied by nearly two orders of magnitude in average synapse count (**Figure 2A**, includes both pre- and postsynaptic sites). For example, 35 BM-Taste neurons innervating Taste bristles on the proboscis accounted for 45% of all head BMN synapses, with an average of 1,028 synapses per neuron. In contrast, 405 eye BM-InOm neurons innervating the interommatidial bristles on the eyes contributed only 9% to the total synapse count due to having the lowest average, with 18 synapses per neuron. Posterior head BMNs had some of the fewest total and average synapses among the BMN types (**Figure 2B,C**, BM-dPoOr, -dOcci, and -vOcci/vPoOr neurons). Taken together, our results show how bristle locations on the head are differentially represented in the brain based on their BMN synapse distributions into distinct zones, and their synapse numbers.

**Figure 2.**
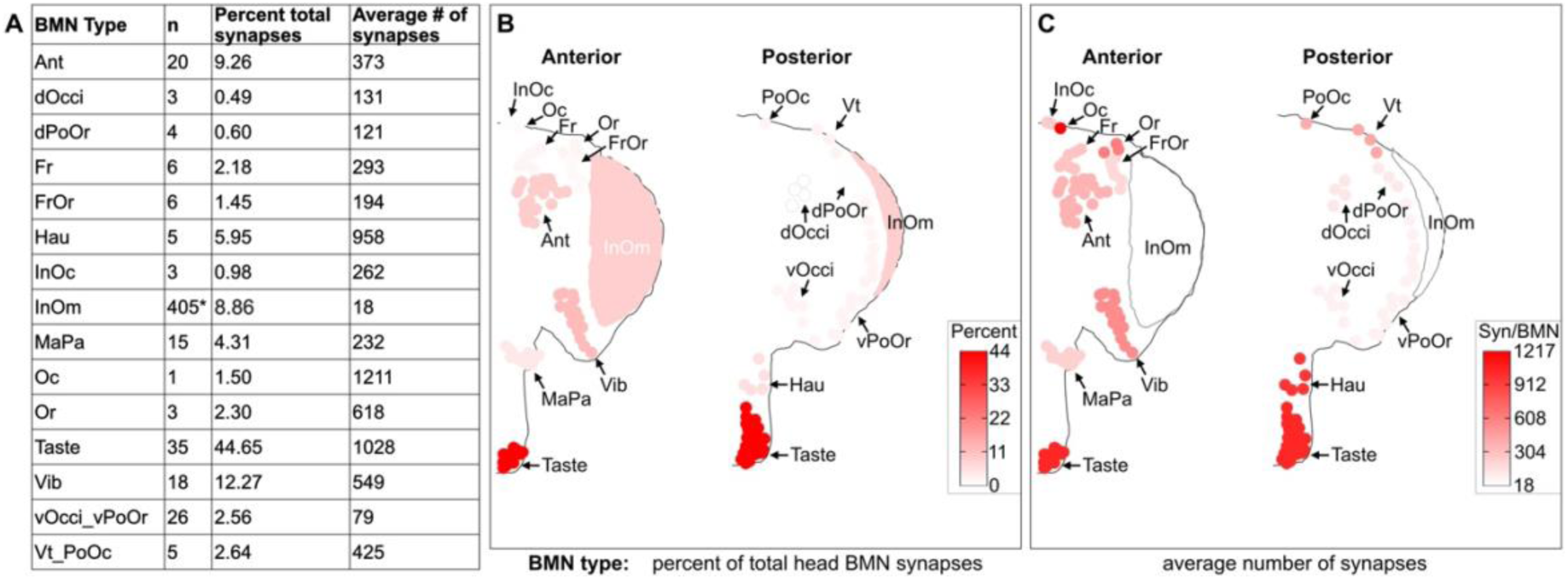
Head BMN type synapse numbers plotted onto their corresponding head bristles. (**A**) Table indicating for each BMN type: numbers of BMNs (n), percent of total head BMN synapses, and the average number of synapses. The percent of total number of synapses was calculated using the total number of input and output synapses for each type, divided by the total synapses for all BMN types. The average number of synapses was calculated using the sum of input and output synapses for each type, divided by the number of BMNs in the type. Box plots of BMN input and output synapses by type are shown in Figure 2 **– figure supplement 1**. (**B-C**) Dot shading on the anterior and posterior head indicates the percent total head BMN input and output synapses (**B**) or the average synapses per BMN type (**C**). *Calculations were done using only BMN synapses from connections to partners that were pre- or postsynaptic by at least five synapses. Therefore, fewer than the total number of BM-InOm neurons were included in the analysis because they did not meet this threshold. Underlying data is in **Supplementary file 1**.

In addition to differing in total synapse number, BMN types vary in their pre- versus postsynaptic composition: all BMNs contain both (Eichler et al., 2024), with presynaptic sites outnumbering postsynaptic sites by ∼2× to ∼9× across types (mean ≈5:1 output-to-input ratio, **Figure 2 – figure supplement 1A,B**, **Supplementary file 2**, **Supplementary file 3**). As expected for sensory afferents, BMNs provide synaptic output to downstream circuits; however, the presence of postsynaptic sites may be less intuitive, and reflects that BMNs can also receive synaptic input onto their central axons within the CNS. These pre- and postsynaptic sites were intermixed within projection zones, showing no clear input/output compartmentalization (**Figure 2 – figure supplement 1C-F**). Codex-rendered individual BMNs also revealed intermixed pre- and postsynaptic site distributions along their axons (Matsliah et al., 2023).

Together, these results show that BMNs project into somatotopically organized zones with intermixed pre-and postsynaptic sites and substantial variation in synapse number and presynaptic-to-postsynaptic ratios, providing a substrate for parallel sensory processing within the grooming circuitry. Notably, if grooming order were driven simply by relative sensory drive—i.e., by BMN types with the strongest synaptic output eliciting cleaning of their corresponding locations first—then synapse number should track the grooming sequence. Instead, differences in synapse number do not align with the order of the grooming sequence: BM-Taste neurons account for the majority of BMN output, yet proboscis grooming is not the first head grooming movement performed, whereas BM-InOm neurons contribute only a small fraction of output despite eye grooming occurring first (**Figure 1E**, **Figure 2A,B**). This indicates that global synapse number alone is not a reliable predictor of the grooming sequence.

### The BMN connectome

The reconstruction and mapping of the head BMNs provided the framework for us to next determine how they interface with pre- and postsynaptic partners. The entire BMN pre- and postsynaptic connectome was previously edited in FAFB by our group and others using the FlyWire.ai platform (Dorkenwald et al., 2024, 2022; Eichler et al., 2024). We focused here on partners of BMNs from the left side of the head. 484 partners that met a 5 synapse connection threshold with BMNs were identified using the FAFB analysis platform Codex (Matsliah et al., 2023). The partners included neurons on both the ipsilateral and contralateral brain hemispheres. Contralateral connections are possible, in part, because some BMN types have projections that cross the midline (Eichler et al., 2024). There were significantly more postsynaptic than presynaptic partners, in agreement with the BMNs containing more presynaptic than postsynaptic structures (**Figure 3A,B**, **Figure 2 – figure supplement 1A,B**, **Supplementary file 2**, **Supplementary file 3, Supplementary file 6**).

**Figure 3.**
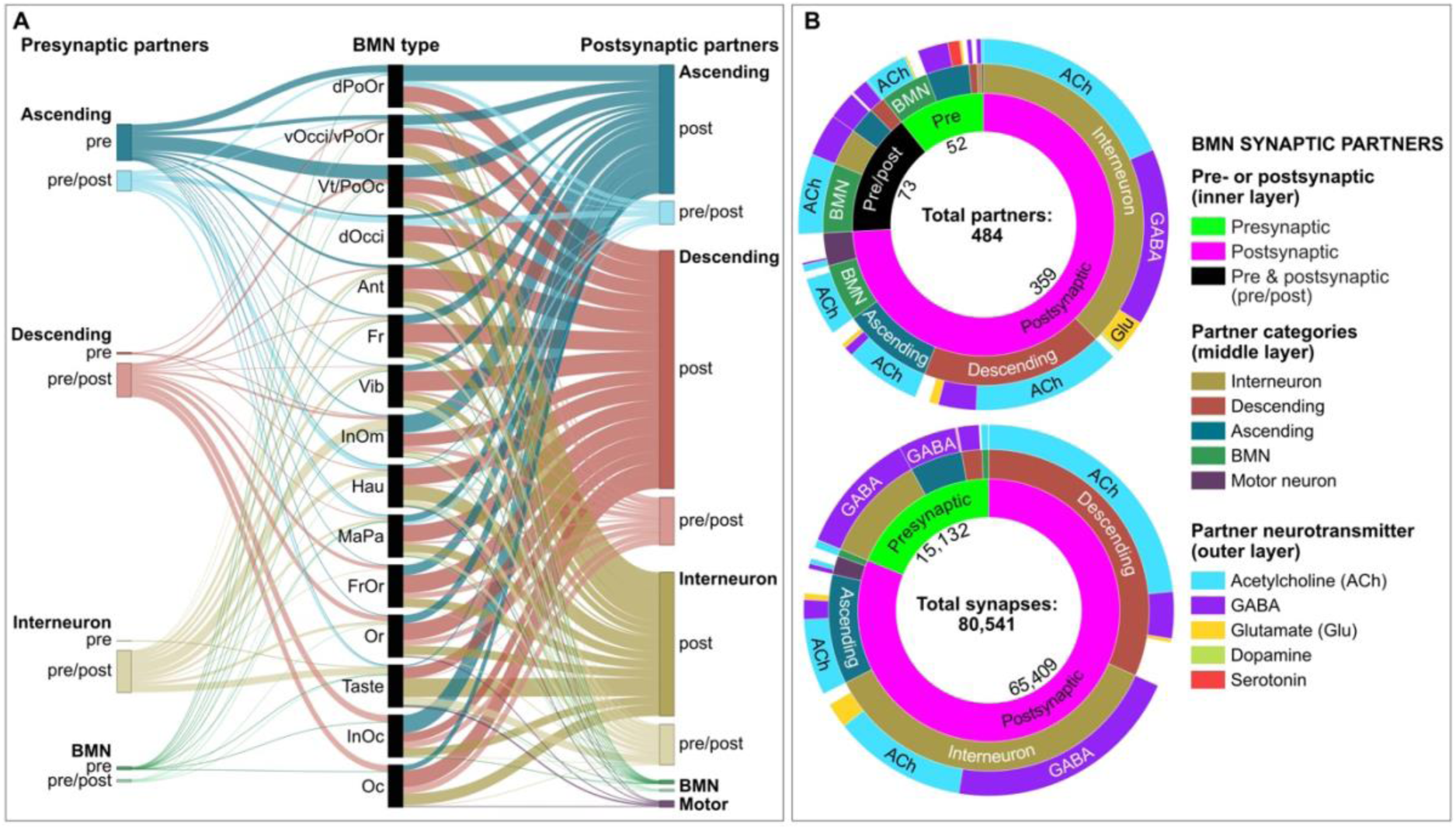
Pre- and postsynaptic connectome of the BMNs. (**A**) Sankey diagram of normalized synapse fractions between BMNs and different partner categories. Black bars in the center of the plot represent the normalized total of postsynaptic sites for each given BMN type, and colored bars along the periphery represent partner categories. All synapse fractions were normalized for visualization by making them proportional to the total number of output synapses for the given BMN type. Boxes to the left of BMNs are presynaptic and those to the right are postsynaptic. Each partner category (except motor neurons) has one subset that is purely pre- or postsynaptic, and another that is both pre- and postsynaptic (pre/post). These subsets are separated and displayed on both sides of BMNs to reveal what proportion of presynaptic input arises from postsynaptic partners. Colors for **A** match the partner categories in **B**. Raw and normalized underlying data are in **Supplementary file 4** and **Supplementary file 5**, respectively. (**B**) Sunburst plots showing the composition of partners (top) or synapses (bottom) that are pre- and postsynaptic to BMNs at a 5 synapse connection threshold. Inner layer shows the proportion of partners or synapses that are presynaptic, postsynaptic, or pre- and postsynaptic (pre- and postsynaptic synapses (bottom plot) include pre/post neurons). Middle layer categorizes the partners or associated synapses as being interneurons, descending or ascending neurons, motor neurons, or BMNs. Outer layer categorizes the partners or synapses based on neurotransmitter prediction. White in the outer rings indicate neurotransmitters could not be predicted. Underlying data are in **Supplementary file 6**.

Partners were grouped into five morphological categories—interneurons, descending neurons, ascending neurons, BMNs, and motor neurons—following FlyWire annotations (Dorkenwald et al., 2024). Interneurons were defined as neurons whose soma and all neurites were confined to the brain. Descending neurons were defined as neurons whose somata are located in the CNS and whose neurites extend into the descending tracts toward the ventral nerve cord (VNC). Conversely, ascending neurons were identified as neurons whose neurites enter the brain through the cervical connective and whose somata lie outside the FAFB imaged volume, resulting in only their neurites being visible in the dataset.

Thus, the BMNs interface with a diverse set of partners whose neurites reside both in the brain and, for ascending/descending classes, extend into the VNC. Because the FAFB dataset includes only the brain and excludes the VNC, the ascending and descending projections outside the brain are not present in the dataset. In addition, because our analysis was restricted to BMNs entering the left hemisphere, the complete right-side BMN connectome is not included, limiting assessment of bilateral symmetry, inter-hemispheric coordination, and variability across sides.

### BMN connectome general organizational features

We identified key organizational features of the BMN pre- and postsynaptic partners in FAFB, based on their connectivity, neurotransmitter identities, and neuron categories (**Figure 3A,B**) (Eckstein et al., 2024; Matsliah et al., 2023). The motor neurons comprise 3% of total BMN partners and were exclusively postsynaptic targets, receiving input from six distinct BMN types and accounting for 2% of total BMN output sites. Subsets of BMNs, ascending neurons, descending neurons, and interneurons are presynaptic, postsynaptic, or both pre- and postsynaptic (pre/post neurons) to the BMNs.

The BMNs are synaptically connected with each other via axo-axonal connections, with BMNs innervating the same or neighboring head bristle populations showing the highest interconnectivity (Eichler et al., 2024). BMNs are likely cholinergic based on neurotransmitter predictions (**Figure 3B**, **Supplementary file 6**) and experimental data (Tuthill and Wilson, 2016b), suggesting somatotopy-based mutual excitation.

Although 18% of BMN synaptic partners are other BMNs, BMN/BMN connections accounted for only 1% of BMN synaptic sites. The majority of synapses were formed with ascending, descending, and interneurons in the CNS. Interestingly, some BMN types lacked specific presynaptic partners from these categories (**Figure 3A**). For instance, BM-InOc and -Oc neurons connect presynaptically with descending neurons but not ascending or interneurons. Conversely, dPoOr neurons on the posterior head connect with ascending presynaptic partners but lack interneuron or descending connections. Thus, while BMN types generally connect with diverse neuron categories, some have more restricted presynaptic inputs.

A striking feature of the presynaptic connectome was that most BMN synaptic input was predicted to be GABAergic (**Figure 3B**, bottom), highlighting a major role for presynaptic inhibition of the BMNs. Further, some of these inputs were from pre/post neurons, revealing feedback inhibition. In contrast to the presynaptic partners, BMN postsynaptic partners mediate both excitation and inhibition downstream, with cholinergic, GABAergic, and glutamatergic partners (**Figure 3B**).

These findings reveal that BMNs engage with diverse partners to form key circuit features, including direct BMN-to-motor-neuron output, mutual excitation among BMNs, presynaptic inhibition, feedback inhibition, and excitatory and inhibitory postsynaptic output. Below, we describe these fundamental features of mechanosensation, and examine how they contribute to grooming behavior.

### BMN postsynaptic motor neurons

Sixteen motor neurons postsynaptic to the BMNs were identified, revealing a direct link between mechanosensory input and motor output (**Figure 4A,B**). These motor neurons were classified by their axonal projections through the labial, pharyngeal, or antennal nerves. Labial and pharyngeal nerve motor neurons produce proboscis movements (McKellar et al., 2020) and connect with BMN types innervating bristles on the proboscis (**Figure 4C-E**, **Supplementary file 7**). Labial motor neurons were exclusively postsynaptic to BM-Taste neurons on the distal proboscis, while pharyngeal motor neurons were connected with BM-Taste and - Hau neuron types innervating neighboring proboscis bristles. Optogenetic activation of BM-Taste neurons elicits proboscis grooming, during which the proboscis often extends (Eichler et al., 2024), suggesting that proboscis extension could be modulated via direct BM-Taste to motor neuron connections. The reconstructed antennal nerve motor neurons likely contribute to antennal movements (Özdil et al., 2024; Suver et al., 2023). These neurons received most of their BMN inputs from BM-Ant neurons on the antennae, but also from BMN types innervating bristles neighboring the antennae (**Figure 4C,F**). Thus, BMNs innervating bristles on the proboscis and antennae connect with motor neurons projecting to those appendages, potentially influencing grooming-related movements. However, BMN inputs accounted for only a small fraction of total synapses onto each motor neuron (≦6.28% of total inputs/BMN type, **Figure 4 – figure supplement 1**, **Supplementary file 7**), suggesting a modulatory contribution rather than direct sensory-driven motor activation.

**Figure 4.**
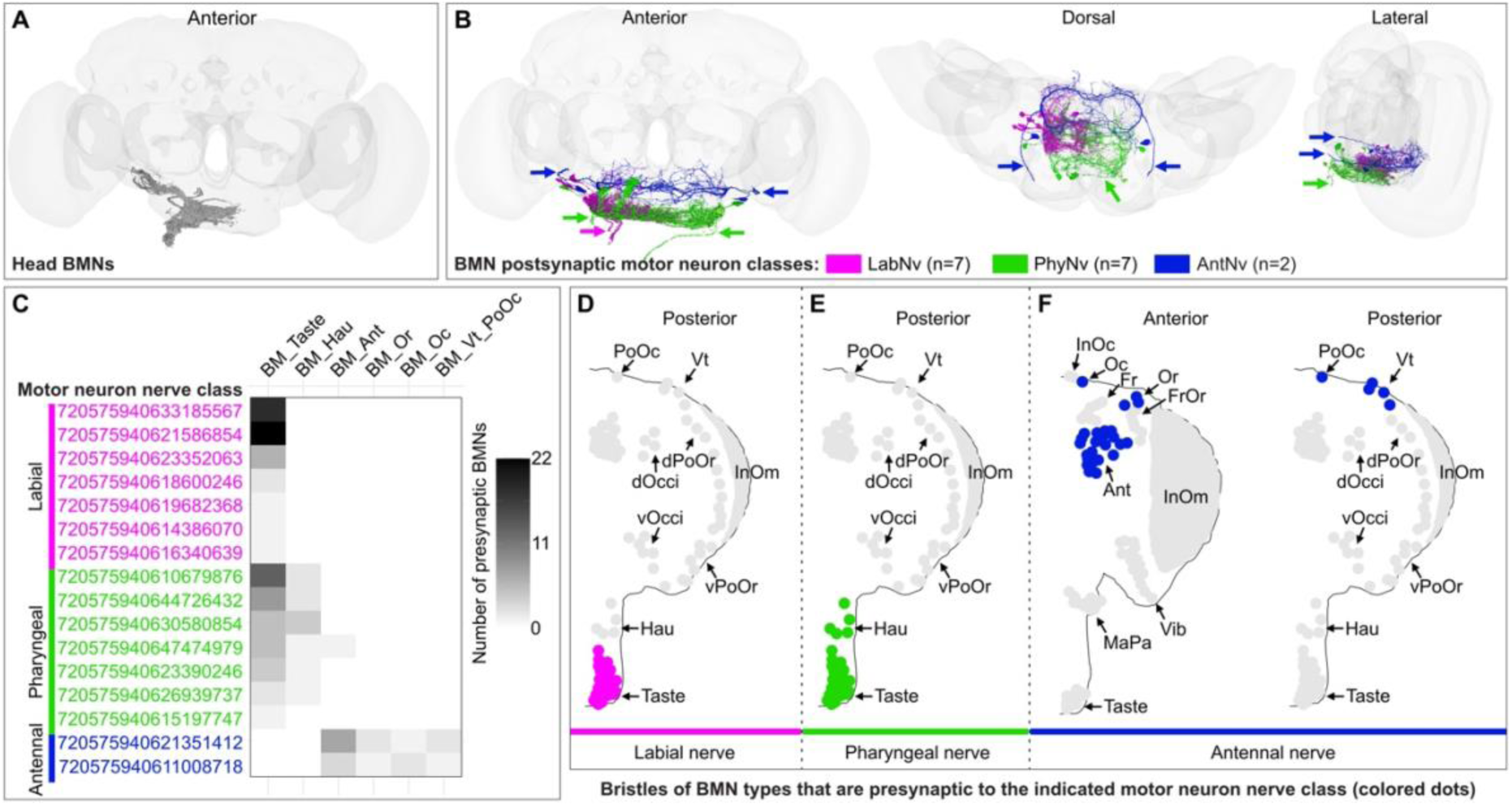
Motor neurons projecting to the proboscis or antennae are postsynaptic to the BMNs on the same appendages. (**A**) Anterior view of reconstructed BMNs projecting into the brain from the anatomical left side of the head. Note: although the BMNs are from the left side of the head, they are displayed in the right brain hemisphere to be consistent with how they are displayed in FlyWire.ai (Schlegel et al., 2024). (**B**) Anterior, dorsal, and lateral views of reconstructed motor neurons that are postsynaptic partners of the BMNs with a 5 synapse connection threshold. Colors indicate the nerve that the motor neuron axons project through to the periphery, including the labial (LabNv, magenta), pharyngeal (PhyNv, green), and antennal (AntNv, blue) nerves. Nerves in both brain hemispheres are indicated with colored arrows. Note: to maintain consistency with FlyWire.ai, all neurons in this manuscript are displayed in the opposite brain hemisphere (e.g left hemisphere neurons are shown on the right; (Schlegel et al., 2024)). (**C**) Heatmap indicating the number of presynaptic BMNs of each type that are connected with different motor neurons. Grayscale shading scale maximum indicates 22 presynaptic BMNs. Colors indicate nerve class. FlyWire.ai neuron identification numbers are shown for each motor neuron. Underlying data are in **Supplementary file 7**. (**D-F**) Colored dots indicate bristles that are innervated by BMN types presynaptic to the indicated motor neuron nerve class, including the labial (**D**), pharyngeal (**E**), and antennal (**F**) nerve classes. (**D,E**) Posterior views. (**E**) The connections of BM-Ant neurons with one of the seven pharyngeal motor neurons (**C**) is not shown. (**F**) Anterior (left) and posterior (right) views.

### BMN synaptic partners in the CNS: ascending, descending, and interneurons

The BMN synaptic partners in the CNS included morphologically diverse sets of ascending, descending, and interneurons (**Figure 5A-J**). Among the 381 partners, most were postsynaptic neurons: 315 postsynaptic, 39 pre/post, and 27 presynaptic (**Figure 5A-D, Supplementary file 6**). Notably, 70% of neurons presynaptic to BMNs were ascending or descending neurons from the VNC (**Figure 5E,G**). The remaining 30% comprised brain interneurons, including exclusively presynaptic (3%) and reciprocal pre/post (27%) neurons, highlighting their role in feedback processing (**Figure 5I**). Postsynaptic connections were predominantly interneurons (56%), with significant contributions from descending (28%) and ascending (16%) neurons (**Figure 5D,F,H,J**). Interneurons are more numerous as distinct partner neurons, whereas descending neurons receive a larger fraction of BMN output synapses across BMN types (**Figure 3A,B**). Thus, descending neurons are fewer in number but tend to receive more BMN synapses per neuron on average, while interneurons are more numerous but often receive fewer synapses per neuron. Together, these partner categories underscore the strong integration of BMNs with local brain circuitry (interneurons), and with pathways linking the brain and ventral nerve cord (VNC), through ascending neurons that provide VNC-derived synaptic input and descending neurons that carry BMN output toward the VNC.

**Figure 5.**
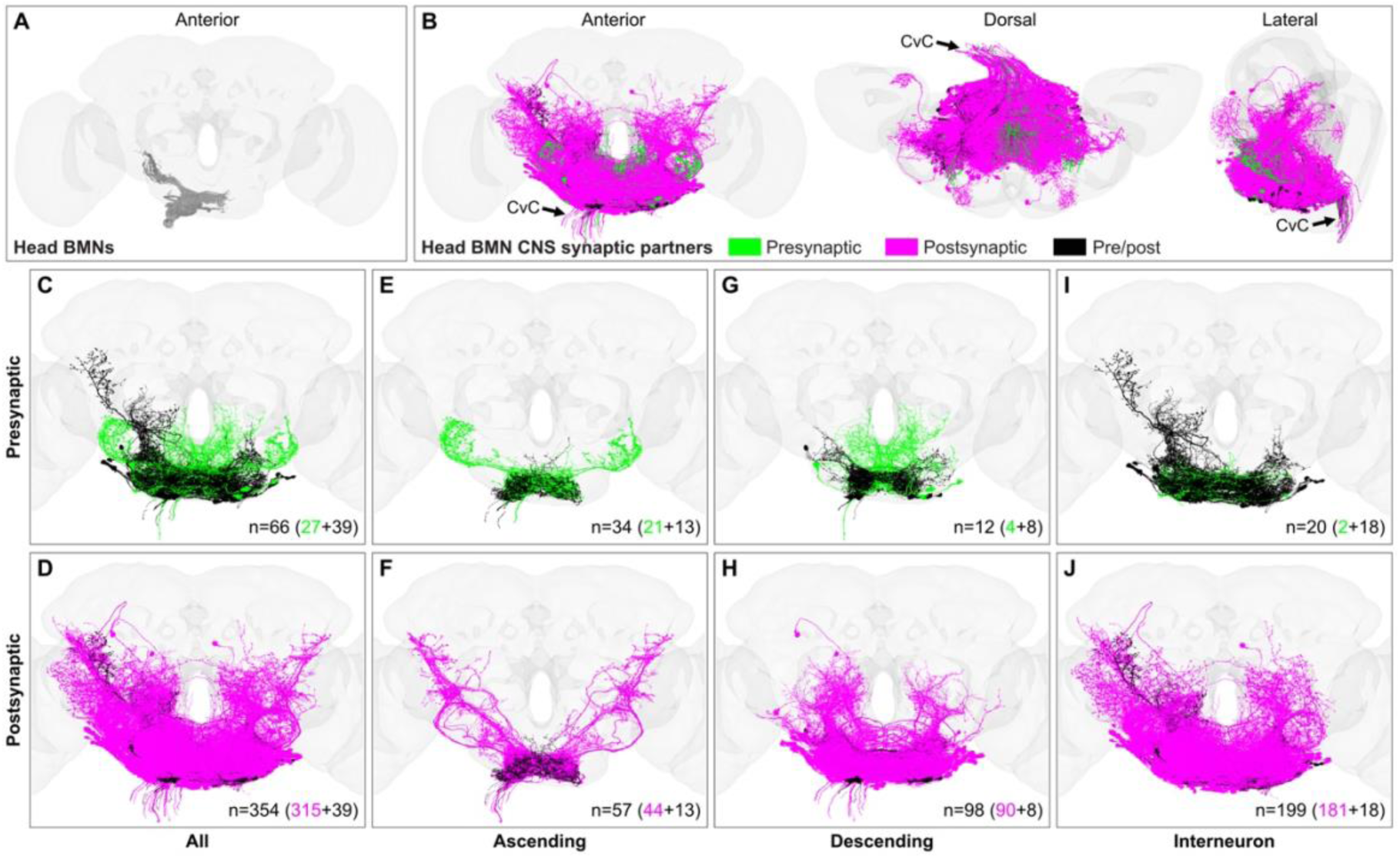
BMN pre- and postsynaptic partners in the CNS. (**A**) Anterior view of reconstructed BMNs projecting into the brain from the anatomical left side of the head. Note: to maintain consistency with FlyWire.ai, all neurons in this manuscript are displayed in the opposite brain hemisphere (e.g left hemisphere neurons are shown on the right; (Schlegel et al., 2024)). (**B**) Anterior, dorsal, and lateral views of reconstructed synaptic partners of the BMNs with a 5 synapse connection threshold (BMN and motor neuron partners not shown). Colors indicate whether the partners are presynaptic (green), postsynaptic (magenta), or pre/post (black) to the BMNs. Cervical connective (CvC) is indicated and contains descending and ascending neuron axons to and from the VNC, respectively. (**C-J**) Anterior view of all pre- (**C,E,G,I**) and postsynaptic (**D,F,H,J**) partners of BMNs. Different categories of synaptic partners are shown in columns, including all partners (**C,D**) ascending (**E,F**), descending (**G,H**), and interneurons (**I,J**). The numbers of the different types are indicated in each panel. Underlying data are in **Supplementary file 6.**

### Synapses of BMN partners are mostly concentrated in the ventral brain

Most BMN partner neurites were concentrated in the ventral brain, with some extending into dorsal regions **Figure 5A-J**). Synapse mapping of nearly all the pre- and postsynaptic sites of BMN partners revealed that partner synapses were predominantly located in the gnathal ganglia (GNG) (**Figure 6A-D**, **Supplementary file 8**, **Supplementary file 9**). The GNG is the ventral-most brain neuropil containing the BMN projections, and connects to the VNC via ascending and descending neurons through the cervical connective (Ito et al., 2014). The GNG also contains interneurons and descending neurons whose activation elicits head grooming movements (Guo et al., 2022; Hampel et al., 2015). On average, BMN partners had 24 times more synapses in the GNG than in their second-highest synapse location, the saddle (SAD), located just dorsal to the GNG. While most synapses were in the GNG and SAD, smaller numbers were found in other neuropils. For instance, presynaptic partners had synapses in the antennal mechanosensory and motor center (AMMC), and in the dorsal and the medial neuropils, vest (VES) and flange (FLA) (**Figure 6A**). Postsynaptic and pre/post partners also had synapses in these regions, along with the wedge (WED) and anterior and posterior ventrolateral protocerebrum (AVLP and PVLP) (**Figure 6B-D**). Some of these neuropils were previously linked to mechanosensory regulation and processing, which may have important implications for BMN-mediated processing (Pacheco et al., 2021; Patella and Wilson, 2018; Suver et al., 2019; Tsubouchi et al., 2017). Thus, while most neuropils containing synapses of second-order BMN partners are located below the esophagus (in the subesophageal zone, SEZ), we found more limited involvement of neuropils in the supraesophageal zone (SPZ; above the esophagus), suggesting relatively limited direct top-down control.

**Figure 6.**
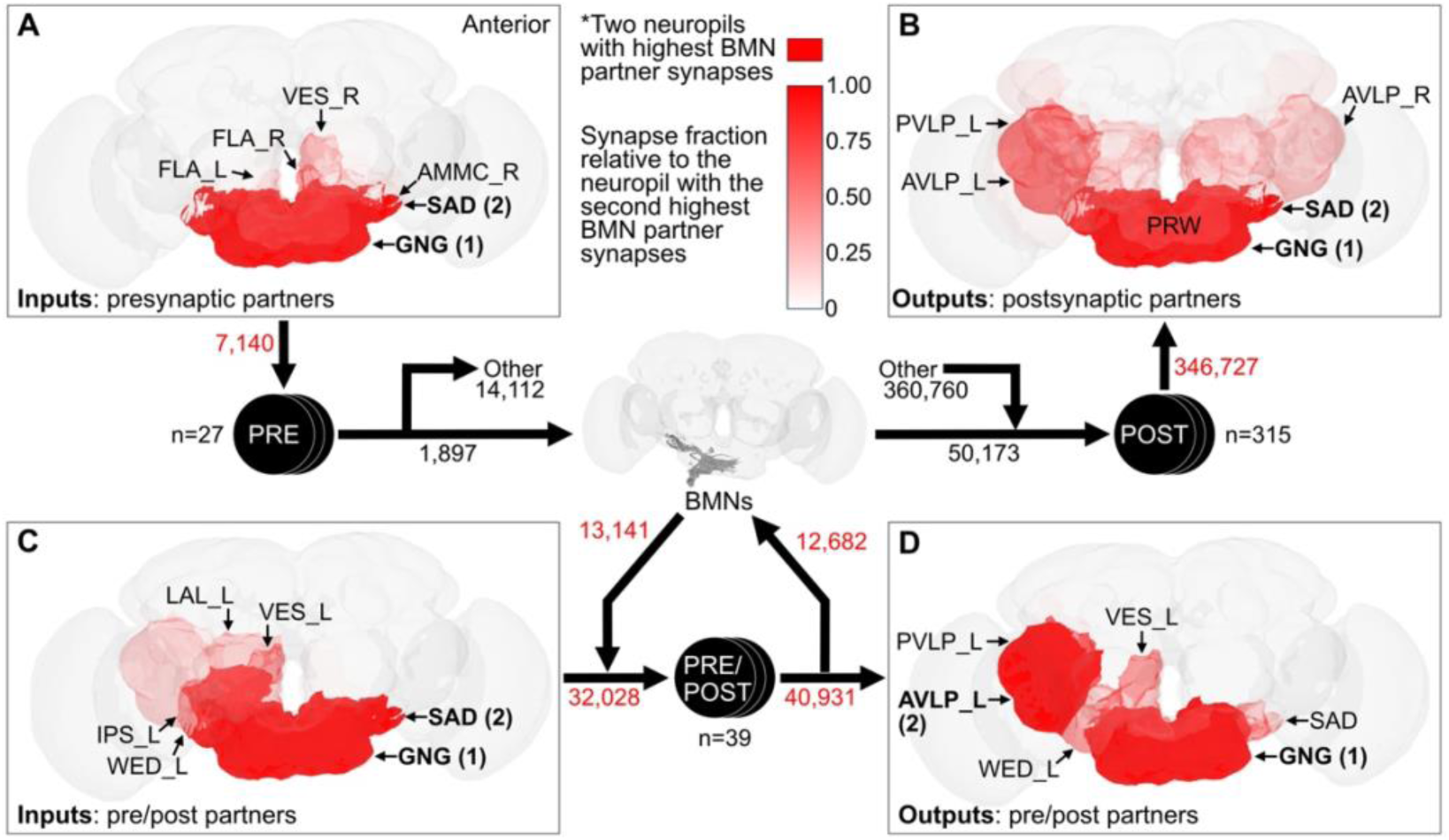
Most BMN partner input and output synapses are distributed in the ventral brain. (**A-D**) Brain neuropil distributions of input or output synapses of the presynaptic (**A**), postsynaptic (**B**), and pre/post (**C,D**) partner neurons. (**A**) Input synapses onto the BMN presynaptic partners. (**B**) Output synapses of the postsynaptic partners. (**C,D**) Input (**C**) and output (**D**) synapses of the pre/post partners. The total numbers of synapses contributing to each panel are indicated in the connectivity diagram in red. For pre/post partners, the number associated with the arrow from the BMNs indicates the BMN input synapses onto pre/post partners, while the arrow to the BMNs indicates pre/post partner output synapses onto BMNs. The other arrows going to and from the pre/post neurons refer to input and output synapses that are not to or from BMNs. Presynaptic partner outputs and postsynaptic partner inputs are not shown, but their synapse numbers with BMNs or other neurons are indicated in black. The 6 neuropils with the most partner synapses are labeled in each panel, with the first and second highest being indicated with bold font. Darker red shades indicate areas with higher synapse counts. Shading is not linear because the gnathal ganglia (GNG) contains most BMN partner synapses. To highlight neuropils with fewer partner synapses, the 2 neuropils with the most synapses are shaded dark red (top shade indicator). All other neuropils are shaded based on the fraction of synapses relative to the neuropil with the second highest number of partner synapses (gradient indicator). Shading is calculated for each panel, and cannot be compared across panels. See **Supplementary file 8** for a complete table of BMN partner pre- and post synapse numbers in all neuropils. Synapses are from ascending, descending, and interneurons, and do include BMNs or motor neurons. Synapses of left side BMN partners shown. Abbreviations: Gnathal ganglia (GNG), saddle (SAD), antennal mechanosensory and motor center (AMMC), vest (VES), flange (FLA), anterior ventrolateral protocerebrum (AVLP), posterior ventrolateral protocerebrum (PVLP), prow (PRW), lateral accessory lobe (LAL), inferior posterior slope (IPS), wedge (WED). Raw and normalized underlying data for partner synapses in the top 31 neuropils (total synapse count > 100) are in **Supplementary file 9**.

### Majority of BMN presynaptic input is GABAergic

GABAergic inhibition is the dominant feature of the presynaptic connectome, as shown by synapse counts from presynaptic or pre/post neurons onto BMNs (**Figure 7A,B**, **Supplementary file 6**). BMNs receive minor inputs from cholinergic BMNs and serotonergic ascending neurons, while the GABAergic inputs originate from ascending, descending, and interneurons (**Figure 7C-J**). Thus, both the brain and VNC contribute to BMN presynaptic input. Partners that are exclusively presynaptic to the BMNs account for 13% of the total input, which is mostly from GABAergic ascending neurons (**Figure 7A,D**). Meanwhile, 84% of BMN presynaptic input is from GABAergic pre/post neurons, suggesting that feedback inhibition is the primary function of BMN presynaptic input (**Figure 7B,H-J**). Notably, 68% of feedback inhibition comes from interneurons, revealing strong influence of BMN feedback inhibition from local neurons (**Figure 7B**). Presynaptic inhibition plays a key role in mechanosensory systems (see Discussion), and is hypothesized to provide a sensory gain control mechanism that establishes the grooming behavioral suppression hierarchy (Hampel et al., 2017; Seeds et al., 2014).

**Figure 7.**
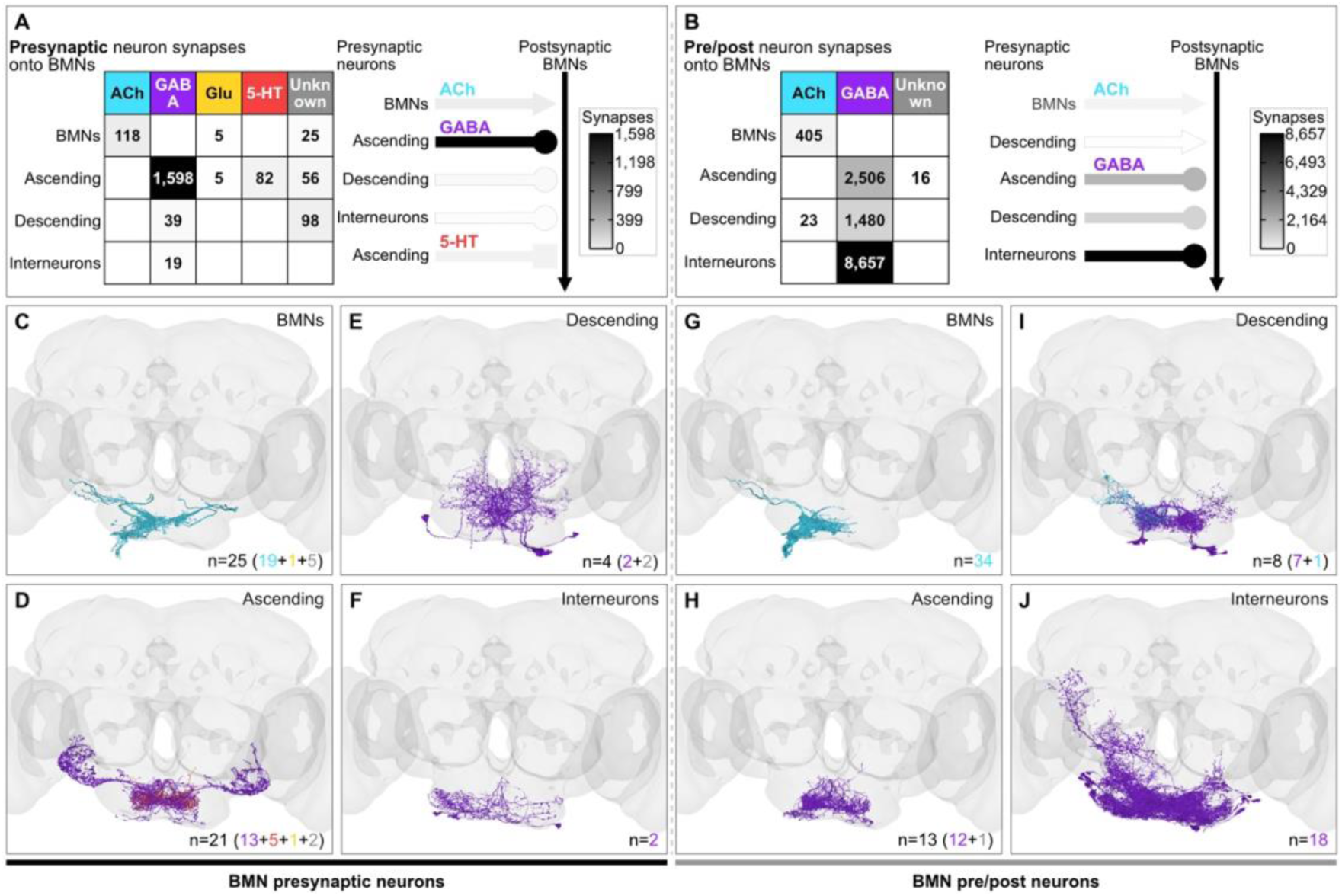
BMN presynaptic inputs are mostly GABAergic. (**A-B**) BMN presynaptic (**A**) and pre/post (**B**) partner neurotransmitters and their numbers of synapses onto BMNs. Table (left) indicates synapses onto BMNs from each partner category. Partners are predicted to use acetylcholine (ACh), GABA, glutamate (Glut), or serotonin (5-HT). Unknown indicates synapses whose neurotransmitter identities could not be determined by the prediction algorithm (Eckstein et al., 2024). Grid grayscale shades indicate synapse numbers, with black indicating 1,598 (**A**) or 8,657 (**B**) synapses (scale indicator on right). Schematic (middle) shows BMN inputs from different presynaptic partners. Edge grayscale indicates the relative numbers of synapses from each category onto BMNs (scale indicator on right). Partner connections are indicated as excitatory ACh (arrow), inhibitory GABA (ball), and modulatory 5-HT (box). Color codes for each neurotransmitter are used in **C-J**. (**C-J**) Anterior views of reconstructed BMN presynaptic (**C-F**) or pre/post (**G-J**) partner categories, including BMNs (**C,G**), ascending (**D,H**), descending (**E,I**), and interneurons (**F,J**). Colors indicate the predicted neurotransmitters for each partner, including GABA (purple), ACh (teal), Glu (mustard), and 5-HT (red). n indicates the total number of neurons in each group. Colored numbers in parentheses represent the number of neurons with predicted neurotransmitter identities, indicated using the color code from panels (**A,B**). See **Supplementary file 6** for a complete table of partners, neurotransmitter predictions, and synapse counts.

### BMN postsynaptic partners are excitatory and inhibitory

Unlike the primarily GABAergic presynaptic partners, the BMN postsynaptic partners include both excitatory and inhibitory neurons (**Figure 8A,B**, **Supplementary file 6**). BMN output synapses were distributed as follows: 53% to excitatory cholinergic neurons, 33% to inhibitory GABAergic neurons, 4% to excitatory/inhibitory glutamatergic neurons, and 10% to neurons that could not be linked to a neurotransmitter by the prediction algorithm (Eckstein et al., 2024; Liu and Wilson, 2013). Notably, 29% of BMN outputs onto exclusively postsynaptic partners targeted cholinergic descending neurons, suggesting a major role for BMNs is to provide feedforward excitation to VNC circuits (**Figure 8A**). BMNs also had significant connections with excitatory and inhibitory interneurons, indicating their integration with local brain circuits to provide both feedforward excitation and inhibition. This excitation is hypothesized in the parallel model to help form BMN feedforward circuits that elicit aimed grooming of specific body locations, while feedforward inhibition could mediate suppression of competing grooming movements (**Figure 1 – figure supplement 1A,B**). Pre/post interneurons provided the majority of GABAergic presynaptic input to BMNs (**Figure 7B**) and also received the largest share of BMN outputs compared with other pre/post neuron categories (i.e., ascending and descending pre/post neurons; **Figure 8B**). These GABAergic interneurons appear to have a major role in BMN feedback inhibition. Taken together, the BMN postsynaptic partners include a diverse set of neurons that mediate both feedforward excitation and inhibition and feedback inhibition, features predicted by the parallel model.

**Figure 8.**
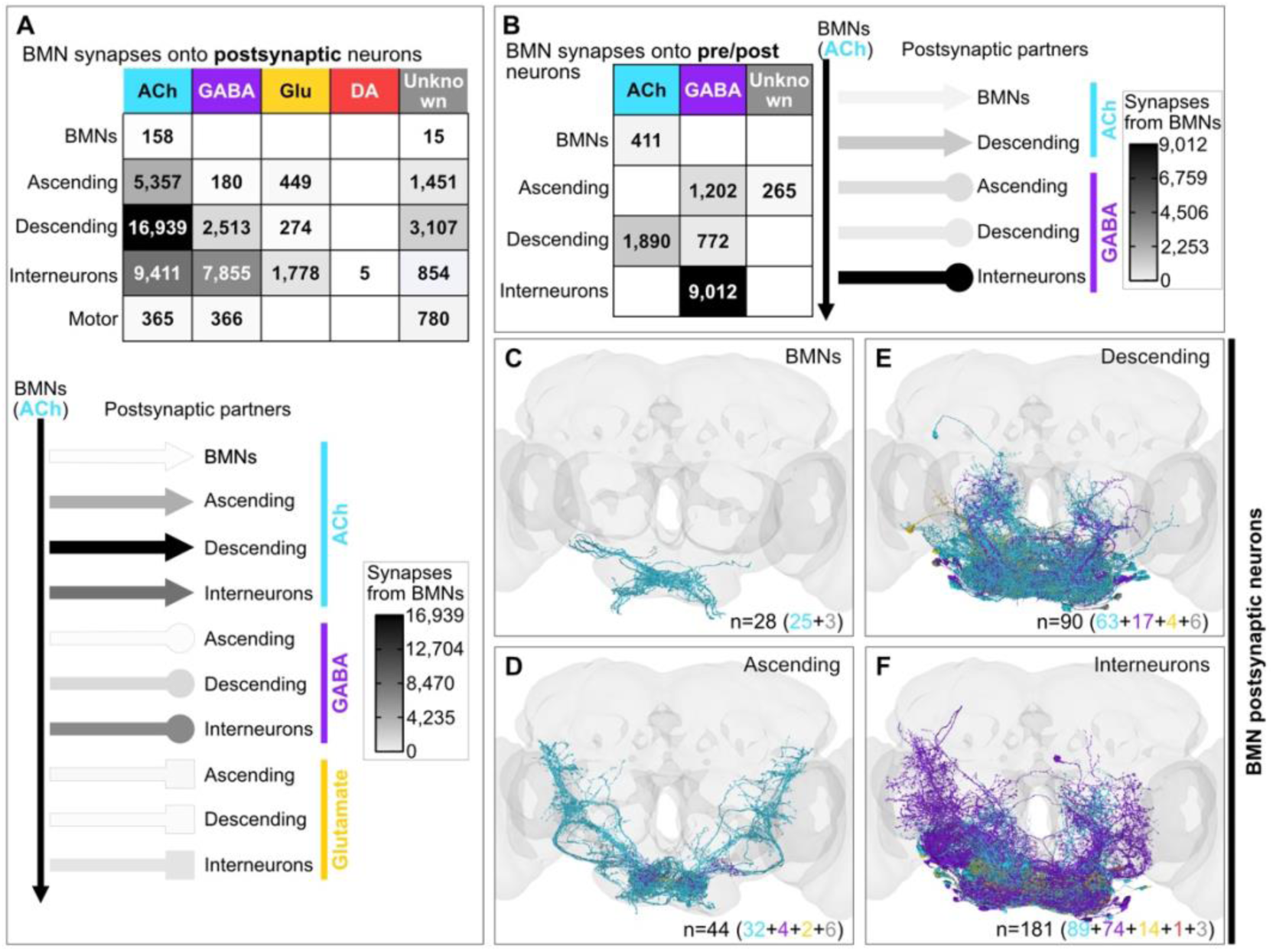
BMN synapses with postsynaptic partners. (**A-B**) BMN synapse numbers onto postsynaptic (**A**) and pre/post (**B**) partners with different neurotransmitter profiles. Table (top or left) indicates the number of BMN synapses onto partners from each partner category. Partners are predicted to use acetylcholine (ACh), GABA, glutamate (Glut), or dopamine (DA). Unknown indicates synapses whose neurotransmitter identities could not be determined by the prediction algorithm (Eckstein et al., 2024). Grid grayscale shades indicate synapse numbers, with black indicating 16,939 (**A**) or 9,012 (**B**) synapses (see scale indicators). Neurotransmitter color codes are used in **C-F**. Schematic illustrates the outputs from BMNs onto postsynaptic partners. Edge grayscale indicates the relative numbers of synapses from BMNs onto each category (see scale indicators). Partners are indicated as excitatory ACh (arrow), inhibitory GABA (ball), and excitatory or /inhibitory Glu (box). DA and motor neurons not shown. (**C-F**) Anterior views of reconstructed BMN postsynaptic partner categories, including BMNs (**C**), ascending (**D**), descending (**E**), and interneurons (**F**). Colors indicate the predicted neurotransmitters for each partner, including GABA (purple), ACh (teal), Glu (mustard), and DA (red). n indicates the total number of neurons in each group. Colored numbers in parentheses represent the number of neurons with predicted neurotransmitter identities, indicated using the color code from panels (**A,B**). See **Supplementary file 6** for a complete table of partners, neurotransmitter predictions, and synapse counts.

### Somatotopy-based connectivity among BMN synaptic partners in the CNS

The head location-specific BMN projection zones shown in **Figure 1A-D** are hypothesized to be where BMN types connect with feedforward excitatory postsynaptic partners in the GNG that elicit aimed grooming of distinct locations (**Figure 1 – figure supplement 1A,C**). We previously found connectomic evidence of such a parallel architecture, showing that BMNs innervating the same or neighboring head bristle populations exhibit similarity in postsynaptic connectivity, whereas those innervating distant bristles demonstrate minimal or no postsynaptic connectivity similarity (Eichler et al., 2024). Here, we extended this analysis to include all BMN pre- and postsynaptic partners in the CNS, and then determined the extent to which the second order connectome reflects the BMN type somatotopic organization.

All head BMNs were first reassessed in pairwise tests for postsynaptic connectivity similarity. As previously reported (Eichler et al., 2024), clustering the BMNs based on cosine similarity scores revealed the highest connectivity similarity among BMNs of the same type (**Figure 9A-C**, **Supplementary file 11**, **Supplementary file 12**). The clustering also revealed different levels of head spatial resolution in connectivity similarity, from the level of BMN types, to the coarser level of BMNs from particular head regions. These different levels of resolution were more apparent when excluding the eye BM-InOm neurons from the analysis. These BMNs are numerous and have relatively low synapse numbers, and therefore obscure cosine similarity clustering among the other BMN types (Eichler et al., 2024). Excluding the BM-InOm neurons from the analysis revealed a coarse level of connectivity similarity among the other BMNs that includes the dorsal, ventral, and posterior head regions (**Figure 9B,D**).

**Figure 9.**
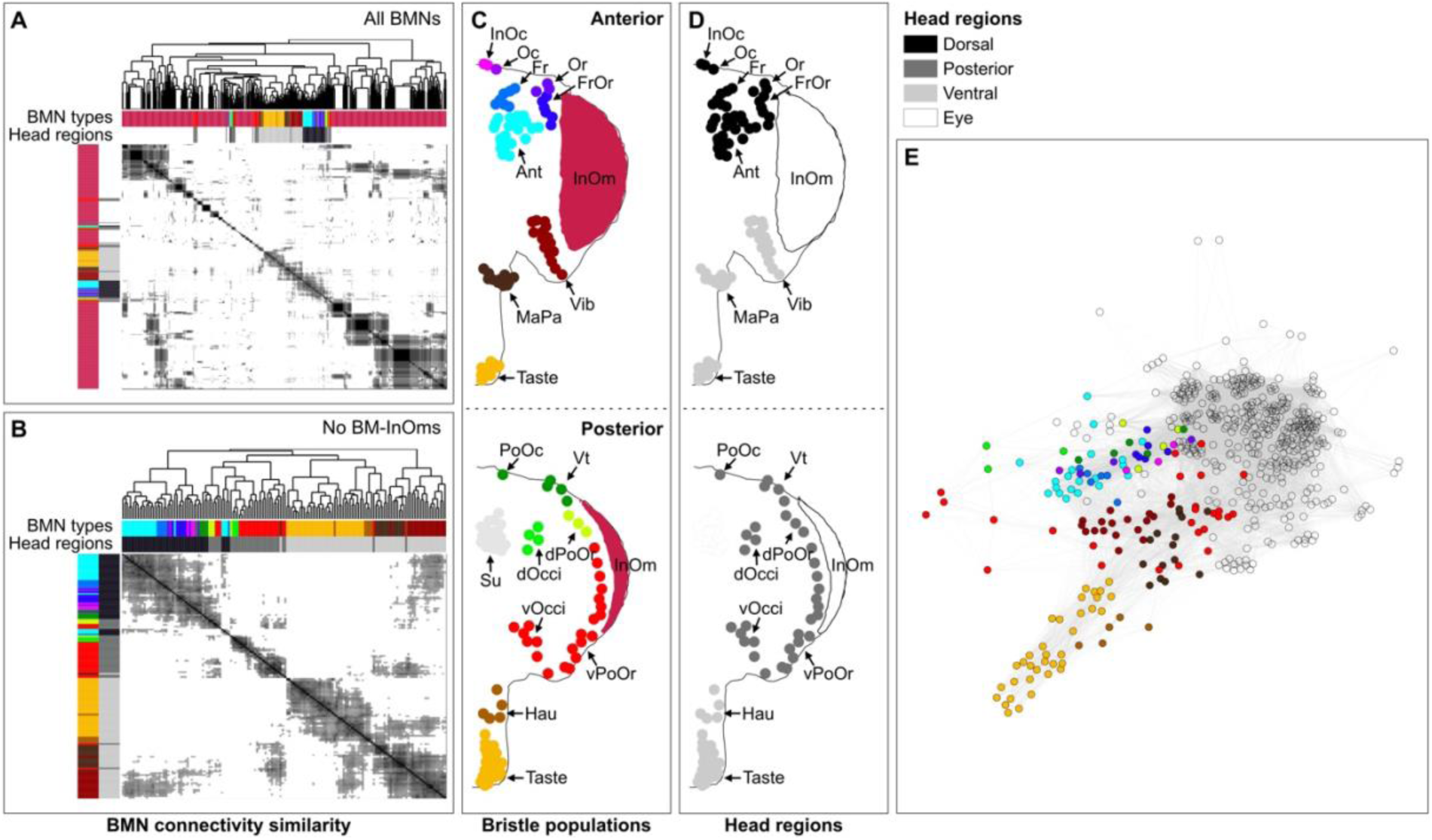
Clustering based on BMN postsynaptic connectivity similarity. (**A,B**) Heatmap of cosine similarity among BMNs based on all output partners in the CNS (ie. ascending, descending, and interneurons; BMN-BMN connections and motor neurons excluded). Row and column order was determined by Ward’s hierarchical clustering method applied to pairwise cosine similarity test scores with a threshold of ≥ 0.3 applied. Clustering of the rows and columns is the same as the dendrogram. Clustering was performed on all BMNs (**A**) and excluding BM-InOm neurons (**B**). Colored bars bordering the headmap indicate the BMN type (**C,** colors) or BMN head region (**D**, grayscale). (**C,D**) Bristle populations on the anterior and posterior head, marked with labeled and color-coded dots indicating their classification by bristle/BMN type (**C**) or head region (**D**). Head regions grayscale code is shown on the top right, and correspond to the dorsal (black), posterior (dark gray), and ventral head (light gray), and eyes (white). (**E**) Simple network diagram of individual BMNs (nodes) based on cosine similarity scores (edges) with a threshold of ≥ 0.3 applied, colored by BMN type. Colored dots indicate BMN type as shown in **C**, with the exception of the BM-InOm neurons that are shown transparent. Node positions were computed with igraph’s default Fruchterman–Reingold force-directed layout: edges (weighted by cosine similarity) act as springs attracting similar neurons, while all nodes repel to avoid overlap. Consequently, neurons with higher similarity scores (i.e., similar postsynaptic connectivity profiles) are drawn close together, forming visually distinct clusters with varying overlap. Underlying data are in **Supplementary file 6**, **Supplementary file 11**, and **Supplementary file 12**.

We next investigated the somatotopy among the pre- and postsynaptic interneurons, descending and ascending neurons by defining their overall connectivity structure with the different BMN types. Because BMNs were clustered by type due to their high connectivity similarity (**Figure 9A-E**), we generated a network graph of their connectome that was based on pooling BMN synapses by type (**Figure 10A-E**, **Supplementary file 10**). Many types had unique pre- and postsynaptic partners, revealing independent BMN-type input and output channels. The BMN types varied widely in their numbers of connected partners, with the BM-Taste neurons on the proboscis having the most partners and the BM-dOcci neurons on the posterior head having the least (**Figure 2 – figure supplement 1B**). Further, BMN types innervating neighboring bristle populations showed common partners, and were in closer proximity in the network graphs than distant BMN types. Three regions with high connectivity similarity included the dorsal, ventral, and posterior head (**Figure 10C,E**, **Figure 9A,B,D**). These results demonstrate somatotopic organization of the BMNs with their pre- and postsynaptic partners at different levels of head spatial resolution, and further reveal the organizational logic of BMN parallel pathways. Thus, the BMNs form largely independent, somatotopically aligned pathways with regional overlap, supporting the idea of parallel grooming circuits.

**Figure 10.**
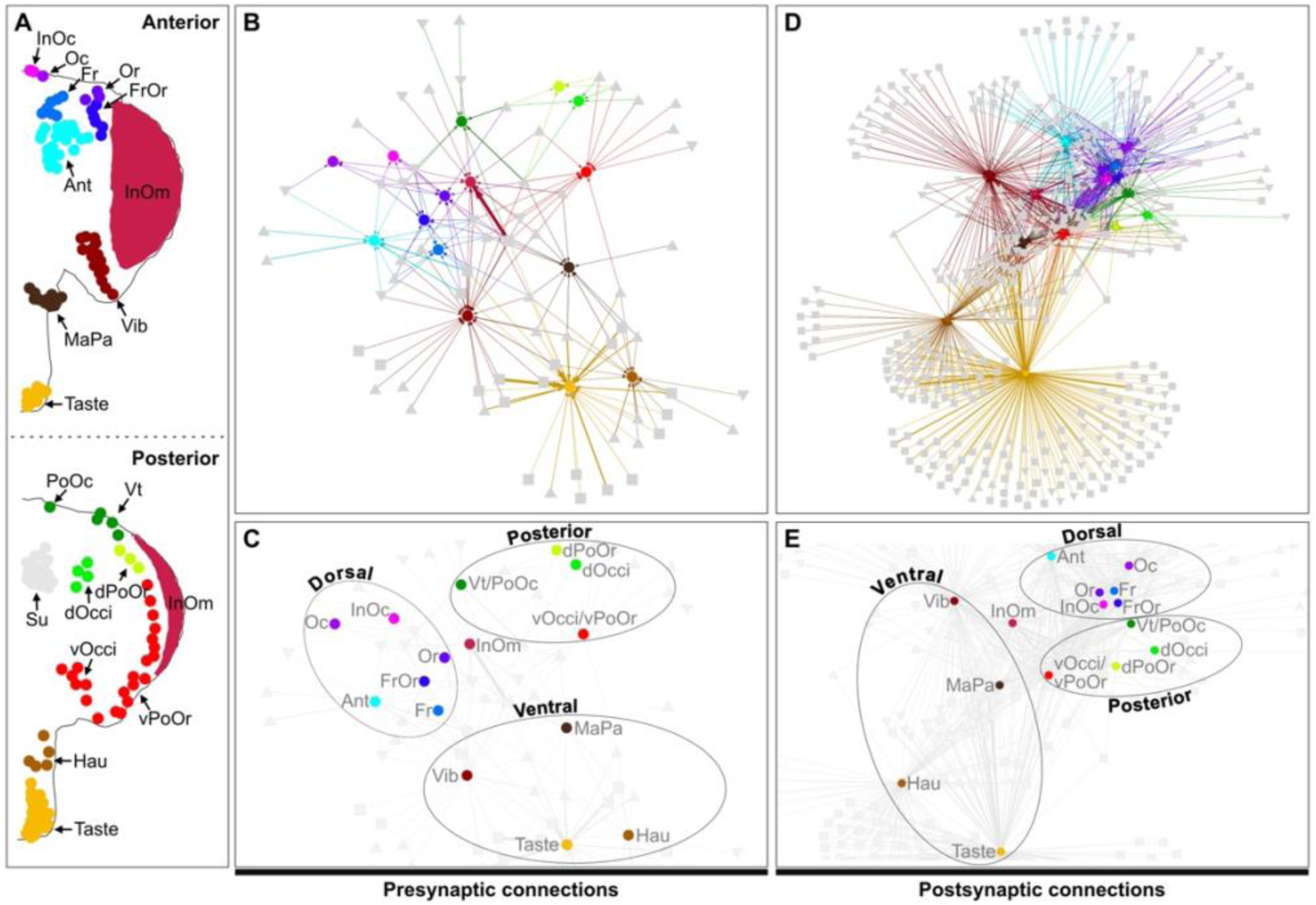
BMN pre- and postsynaptic connectomes show somatotopic organization. (**A**) Bristle populations on the anterior and posterior head, marked with labeled and color-coded dots indicating their classification. Bristle colors used in **B-E** refer to their corresponding BMN types. (**B-E**) Network diagrams of connections between BMN types (colored dots; synapses pooled, each node represents all BMNs of that type) and their presynaptic (**B,C**) or postsynaptic partners (**D,E**) shown in gray. Squares correspond to interneurons while down and up triangles indicate descending and ascending neurons, respectively. Each node represents an individual partner. BMN/BMN connections and motor neurons are excluded. Neurons that are both pre- and postsynaptic are included in all network diagrams in **B-E**. **B** and **D** show the entire network diagram, while **C** and **E** are zoomed cutouts with edge colors removed to better demonstrate position of BMN type nodes relative to each other. (**B,D**) Edges are in colors corresponding to the BMN type inputs (**B**) or outputs (**D**). (**C,E**) Edges are gray. Ellipses indicate BMN types in particular head regions, including the dorsal, posterior, and ventral head (see Figure 9D for graphical view of regions on the head). Node positions were computed using the visNetwork-igraph interface to employ a Fruchterman–Reingold force-directed layout: each directed edge acts like a spring whose strength is proportional to the edge weight (synapse count), drawing highly connected nodes together, while all nodes repel one another to prevent overlap. Consequently, neurons with many or strong synaptic connections cluster closely in the 2D embedding, whereas weakly or unconnected neurons are pushed to the periphery. Underlying data are in **Supplementary file 6** and **Supplementary file 10**.

### BMN feedback inhibition is mediated by somatotopically organized pre/post neurons

The BMN pre- and postsynaptic partners showed somatotopy-based connectivity corresponding to specific head locations (**Figure 10A-E**). Among these partners, 39 were pre/post neurons, and it was unclear to what extent their reciprocal connectivity was somatotopically organized—i.e., whether they primarily form reciprocal loops with the same BMN type, and whether some also link different BMN types by receiving input from one type while providing output to another. It was also unclear whether BMN–pre/post connectivity was grossly asymmetric—serving almost exclusively as presynaptic or postsynaptic targets with only relatively few reciprocal contacts—or instead formed robust bidirectional connections with BMNs. Using a presynaptic:postsynaptic connectivity ratio filtered to 0.1–10, five neurons were excluded for asymmetric wiring and 34 pre/post neurons were used for further analysis.

A network diagram including these 34 pre/post partners showed that all connected with one or more BMN types, but not all connections were reciprocal (**Figure 11 – figure supplement 1**, **Supplementary file 13**). Notably, a subset of these pre/post neurons formed non-reciprocal connections between different BMN types, for example receiving input from eye BM-InOm neurons while providing output to antennal BM-Ant neurons. Such cross-type inhibitory pathways are notable because they could mediate heterotypic suppression between head regions in a direction consistent with the grooming suppression hierarchy (e.g. eye suppresses antennal grooming, **Figure 1 – figure supplement 1B**). However, we also identified cross-type inhibitory pathways that proceeded in directions not consistent with the model of hierarchical suppression.

**Figure 11.**
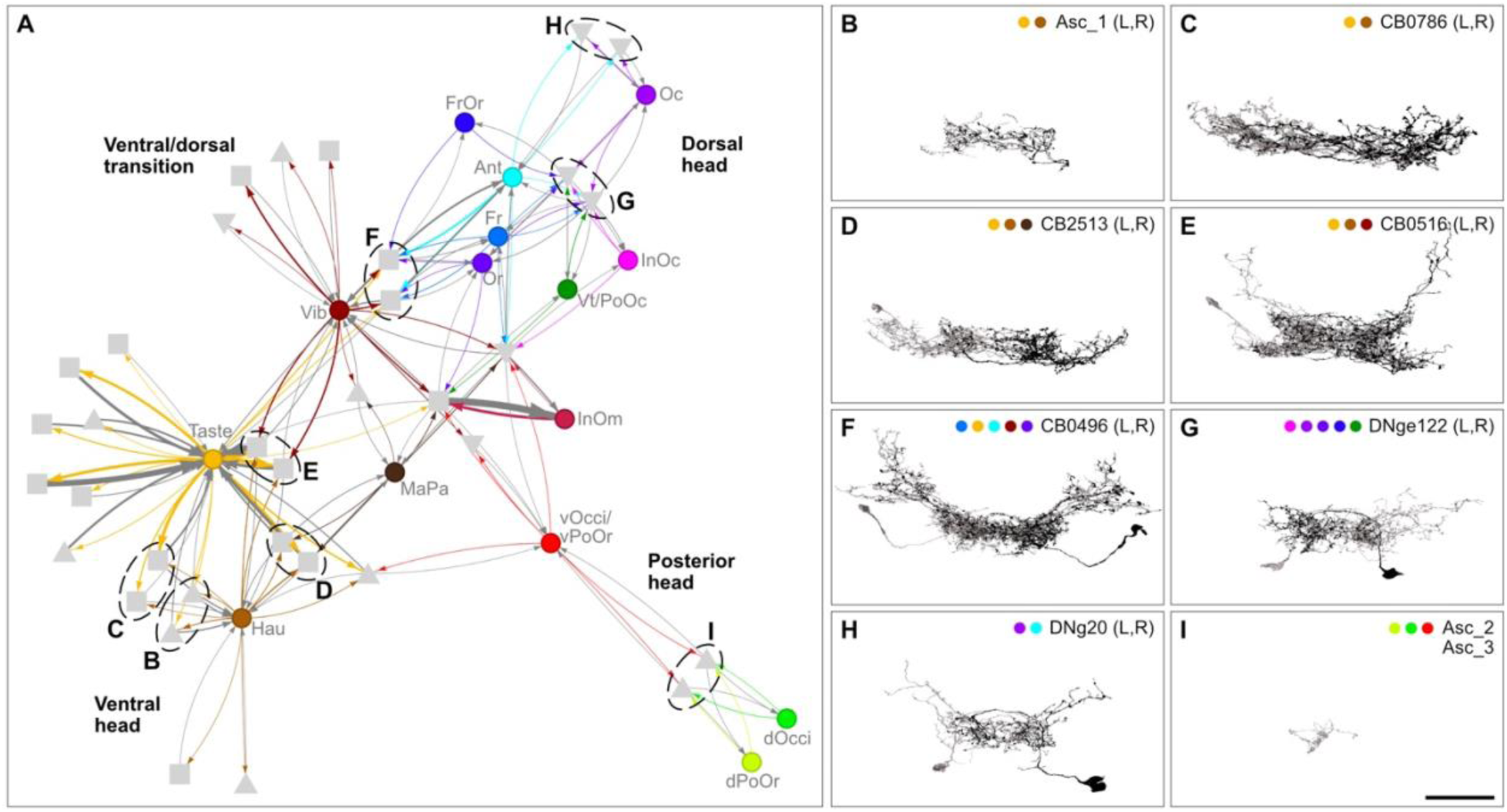
Pre/post neuron pairs form feedback loops with particular BMN types. Connections between BMN types (colored dots) and neurons that are reciprocally pre- and postsynaptic to particular BMN types (gray). Squares correspond to individual interneurons while down and up triangles indicate descending and ascending neurons, respectively. Only edges contributing to reciprocal BMN and pre/post neuron connections are shown. Edges are in colors corresponding to the BMN type outputs, while gray edges correspond to BMN GABAergic inputs. All edges with non-reciprocal connections with pre/post neurons were removed. Bold text indicates the general head location of BMNs on the plot, revealing somatotopy-based connectivity with pre/post neurons (i.e. ventral, dorsal, posterior, and the ventral/dorsal transition). Node positions were computed using a Fruchterman–Reingold force-directed layout: each directed edge acts like a spring whose strength is proportional to the edge weight (synapse count), drawing highly connected nodes together, while all nodes repel one another to prevent overlap. Consequently, neurons with many or strong synaptic connections cluster closely in the 2D embedding, whereas weakly or unconnected neurons are pushed to the periphery. Ellipses indicate neurons shown in **B-I**. See Figure 11 **– figure supplement 1** and **Supplementary file 13** for a complete account of the pre/post neuron connections with the BMN types and information used for pre:post ratio and reciprocal filtering. (**B-I**) Some BMN pre/post neurons include pairs of neurons from the left and right (L,R) brain hemispheres. Shown are pre/post neuron pairs that are connected with specific BMN types, including ascending neurons Asc_1 (L,R) (**B**), interneurons CB0786 (L,R) (**C**), CB2513 (L,R) (**D**), CB0516 (L,R) (**E**), CB0496 (L,R) (**F**), descending neurons DNge122 (L,R) (**G**), DNg20 (L,R) (**H**), and ascending neurons Asc_2 and _3 (**I**). All neurons are of the same cell type (morphologically similar pairs), with the exception of the non morphologically similar ascending neurons shown in **I**. (**B**) Asc_1 (L,R) are not a pair like the others because they are not annotated as such in Codex due to difficulties annotating ascending neurons in the FAFB dataset. However, their axons in the brain are morphologically similar. Colored dots in each panel correspond to BMN types that are connected to both neurons in the pair. Scale bar, 50 μm.

All 34 pre/post neurons formed reciprocal connections with specific BMN types, leading us to separately evaluate these putative local feedback motifs. To more clearly visualize reciprocal connectivity between pre/post partners and each BMN type, we removed non-reciprocal edges for this analysis (**Figure 11A**). Among the reciprocal connections, some were exclusive to particular BMN types (e.g. pre/post neurons dedicated to BM-Taste or -Vib neurons), whereas others reciprocally connected with multiple BMN types from neighboring head locations and tended to group with BMN types from the ventral, dorsal, and posterior head, with some connections involving BMN types positioned near the ventral–dorsal transition (**Figure 11A**).

Together, these results indicate that pre/post connectivity with BMNs is somatotopically organized. Because these pre/post neurons are predicted to be GABAergic, the reciprocal motifs are consistent with local feedback inhibition that can regulate BMN gain within a head region, while the non-reciprocal motifs provide a potential substrate for cross-channel suppression between BMN types (**Figure 1 – figure supplement 1B**).

The network diagram of 34 reciprocally connected pre/post neurons revealed eight distinct neuron pairs that were tightly clustered, with each pair showing striking similarity in their connectivity with specific BMN types (**Figure 11A**, ellipses). Interestingly, while one of these eight pairs showed limited morphological similarity with each other, the other seven were morphologically identical neurons from the left and right brain hemispheres (**Figure 11B–I**). Six of these identical pairs were previously annotated in Codex as present bilaterally (CB0786, CB2513, CB0516, CB0496, DNge122, DNg20). The pairs formed reciprocal connections with multiple BMN types from the same head regions (ventral, dorsal, posterior). This reveals that head region-specific, bilaterally symmetric pre/post neuron types mediate somatotopy-based feedback inhibition onto BMNs.

### Postsynaptic partner developmental origins

We next focused on defining the structure of the parallel BMN postsynaptic excitatory pathways that elicit grooming movements (**Figure 1 – figure supplement 1A,C**), first categorizing the postsynaptic partners based on developmental birth timing. In the *Drosophila* brain, primary neurons derived from embryos function throughout larval and adult stages, while secondary neurons derived post embryonically function exclusively in adults. The FAFB-reconstructed descending neurons and interneurons were previously categorized as putative primary or secondary neurons, with ascending neurons remaining unclassified (Schlegel et al., 2024). Applying these classifications to the BMN postsynaptic partners revealed 163 putative primary neurons and 122 secondary neurons (**Figure 12 – figure supplement 1A-G**, **Supplementary file 14**). Putative primary neurons received 59% of BMN output synapses, including mostly cholinergic descending neurons and GABAergic interneurons (**Figure 12 – figure supplement 1AB,E**, **Figure 12 – figure supplement 2**).

**Figure 12.**
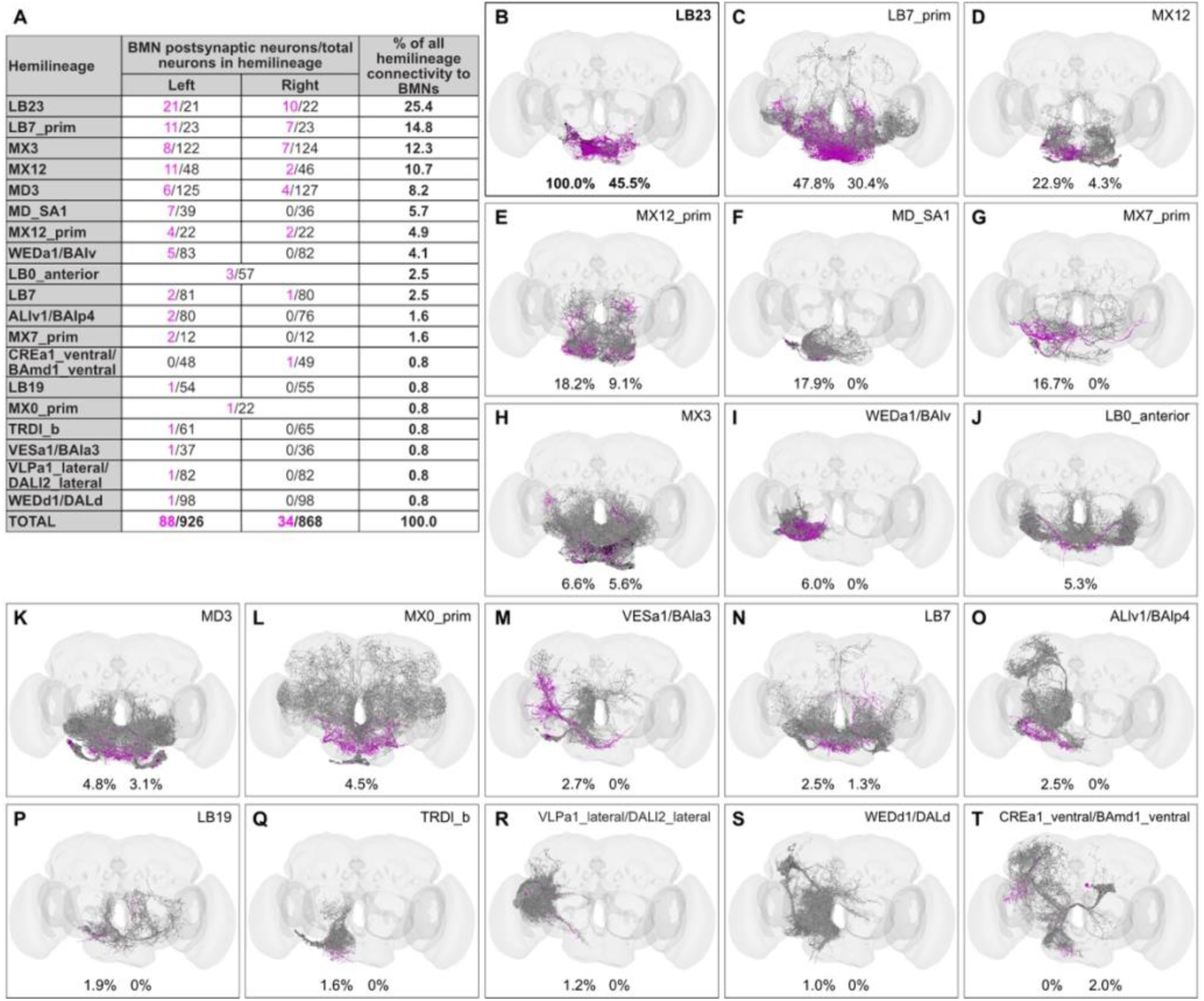
BMN postsynaptic hemilineages. (**A**) Each hemilineage containing neurons that are postsynaptic to left-side BMNs is listed in the first column. In the second column, the number of post and pre/post neurons per hemisphere that are postsynaptic to BMNs is indicated (magenta numerator), along with the total number of neurons in each hemilineage (denominator). Both ipsilateral (left) and contralateral (right) neurons are included. The third column represents the percentage of total BMN-hemilineage connectivity that is accounted for by each hemilineage. This percentage is calculated as the number of neurons in the hemilineage that are connected to BMNs, divided by the total number of BMN-connected hemilineage neurons (122). (**B-T**) Neurons within each hemilineage that are postsynaptic targets of BMNs (magenta) and remaining hemilineage neurons that are not connected with BMNs (grey). The percentages below each hemisphere indicate the proportion of BMN postsynaptic partners within the hemilineage, calculated relative to the total number of neurons in that hemilineage. Information about the BMN postsynaptic partner developmental origins, synapse counts, and predicted neurotransmitters are in Figure 11**—figure supplement 1**. For a list of connected hemilineages and their relative connectivity with all BMN partners, see Figure 11**—figure supplement 2**. Underlying data are in **Supplementary file 14**.

Secondary neurons received 21% of BMN synapses, primarily onto excitatory cholinergic neurons, but also onto some GABAergic neurons (**Figure 12 – figure supplement 1A,C,F**). Taken together, the BMN postsynaptic connectome includes both primary and secondary neurons. The primary neurons form mainly descending excitatory and local inhibitory pathways, whereas the secondary neurons form mainly descending and local excitatory pathways.

### The entire cholinergic hemilineage 23b (LB23) is postsynaptic to BMNs

To identify neurons crucial for establishing the BMN-postsynaptic parallel pathways that elicit head grooming movements, we focused on secondary hemilineages. In the *Drosophila* CNS, a hemilineage refers to the cohort of neurons derived from a single stem cell-like neuroblast that share a common developmental origin, stereotyped morphology, and are thought to have related functional roles within a circuit (Harris et al., 2015; Wreden et al., 2017). This focus was motivated by earlier findings that neurons whose activation elicited head grooming had morphologies consistent with specific hemilineages (Hampel et al., 2015; Seeds et al., 2014). We hypothesized that some of these neurons could be postsynaptic to the BMNs. Nineteen hemilineages were found to be postsynaptic to BMNs, with most having fewer than 25% of their members connected (**Figure 12A-T**, **Figure 12 – figure supplement 2**). Hemilineage 23b on the other hand, referred to here as LB23, stood out with 100% of its members being connected with BMNs on the ipsilateral side and 46% on the contralateral side (**Figure 12B**). The LB23 neurons accounted for 7% of all BMN postsynaptic partners, and received 12% of the total BMN synapses. This high connectivity suggested a critical role for LB23 in BMN mechanosensory processing.

### LB23 hemilineage anatomically corresponds to grooming-related neurons

The LB23 hemilineage is located in the GNG, where neurons involved in head grooming were previously identified. We previously described different transgenic driver lines expressing in subsets of GNG neurons that elicit head grooming movements with optogenetic activation (**Figure 13A**) (Hampel et al., 2015; Seeds et al., 2014). Expression of green fluorescent protein (GFP) in these neurons revealed morphological similarity with the EM reconstructed LB23 neurons. Interestingly, the R40F04-GAL4 driver line, published in Seeds 2014, expresses in neurons whose activation elicited grooming of different locations on the whole head, including the eyes, antenna, ventral and dorsal head. This line labeled a cluster of 14 neurons in the GNG that collectively showed morphological similarity with the projections of the entire reconstructed LB23 hemilineage of 21 neurons (**Figure 13B,C**, arrows, **Supplementary file 15**). However, R40F04-GAL4 also expressed in dorsal brain neurons, raising the possibility that these neurons elicit head grooming.

**Figure 13.**
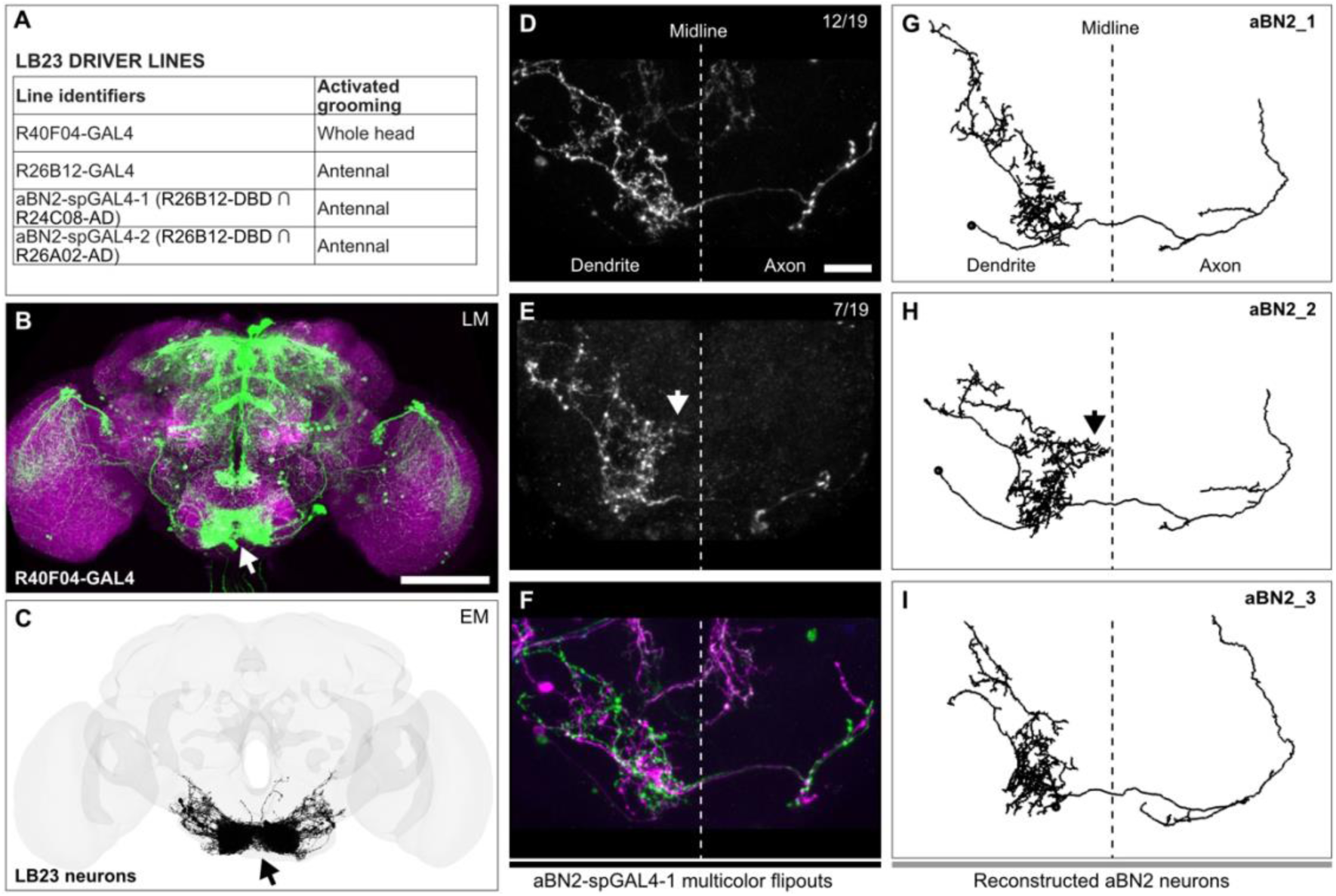
Hemilineage 23b (LB23) neurons elicit head grooming movements. (**A**) Published driver lines expressing in neurons whose activation elicit head grooming (Hampel et al., 2015; Seeds et al., 2014). Shown are the identifiers for each line and their activated head grooming movements. (**B**) Anterior view of a light microscopy (LM)-imaged brain of the R40F04-GAL4 driver line expressing GFP in LB23 neurons (arrow) and other neurons. Image is a confocal Z-stack maximum intensity projection, immunostained for Bruchpilot (magenta) and GFP (green). Scale bar, 100 μm. (**C**) EM reconstructed LB23 neurons from both brain hemispheres (21 neurons on left, 22 on right). (**D,E**) MCFO labeling of aBN2 neurons targeted by aBN2-spGAL4-1. Shown are, maximum intensity projections of two aBN2 neurons stained using tag-specific antibodies (anterior views). The neuron shown in **E** has a midline projecting branch (arrow). Panel top right corner indicates the number of labeled aBN2 neurons without (**D**) or with (**E**) the midline projecting branch, versus the total MCFO-labeled aBN2 neurons. Dotted line indicates the brain midline. Scale bar, 20 μm. (**F**) Labeling of two aBN2 neurons in the same brain hemisphere that both lacked the midline-projecting branch. The green neuron is shown in **D**. (**G-I**) The three left hemisphere EM reconstructed aBN2 neurons (names indicated in top right corner). Dotted line indicates the brain midline. (**H**) Midline projecting branch of aBN2_2 is indicated with an arrow. Neurons rendered from the FAFB reconstructions in CATMAID (Hampel 2020). The following are the flywire IDs for the different aBN2 neurons: aBN2_1 (720575940634683237), aBN2_2 (720575940618961857), aBN2_3 (720575940621355759).

We also identified driver lines expressing in neurons eliciting aimed grooming of the antennae (**Figure 13A**, R26B12-GAL4, aBN2-spGAL4-1 and -2). Prior to being recognised as LB23 neurons, a subset of three were shown to be necessary and sufficient for eliciting antennal grooming and named antennal brain neurons 2 (aBN2) (Hampel et al., 2015). There are three proposed aBN2 neurons based on the observation that aBN2-spGAL4-1 labels three neurons whose silencing disrupts antennal grooming. We used aBN2-spGAL4-1 with the multicolor flipout (MCFO) method to stochastically label each neuron individually and determine the extent to which these three neurons resembled LB23 neurons (Hampel et al., 2020b; Nern et al., 2015). Indeed, the labeled aBN2 neurons were nearly morphologically identical, with an ipsilateral dendritic field and an axon projecting to the contralateral brain hemisphere (**Figure 13D,E**, two examples). 7 out of 19 (37%) labeled aBN2 neurons had a dendritic branch approaching the brain midline (**Figure 13E**, arrowhead). We concluded that one of the three aBN2 neurons has this midline branch, given that two were labeled in the same brain hemisphere that both lacked the branch (**Figure 13F**).

There are 21 LB23 neurons in the FAFB left brain hemisphere (22 on the right) that consist of different morphologically distinct neurons including 18 interneurons and 3 descending neurons (**<u>Flywire.ai link 1</u>**). Among these diverse interneurons, we identified three that resembled the MCFO-labeled aBN2s (**Figure 13G-I**, named aBN2_1, 2, and 3). One of these neurons (aBN2_2) had a midline-projecting dendritic branch, in agreement with what we predicted from the MCFO data (**Figure 13E,H**). These morphological similarities provide strong evidence that aBN2 neurons belong to the LB23 hemilineage. When taken with behavioral evidence (Hampel et al., 2015), it uncovers a role for LB23 hemilineage neurons in eliciting aimed head grooming movements.

### BMNs and LB23 neurons form somatotopic pathways that elicit aimed grooming

In accordance with the parallel model of grooming, we hypothesize that BMNs connect with somatotopically organized excitatory parallel pathways eliciting aimed grooming of specific head locations (**Figure 1 – figure supplement 1A,C**). LB23 neurons could form these pathways, given that they are cholinergic and their activation elicits aimed grooming. We tested the extent to which BMNs and LB23 neurons connect to form a parallel architecture. Indeed, a network graph revealed that subpopulations of LB23 neurons are preferentially connected with particular BMN types from different head bristle locations (**Figure 14A,B**, **Supplementary file 16**). Some LB23 interneuron members connected exclusively with particular BMN types, highlighting bristle population specific LB23 pathways (e.g. BM-Taste and -Hau neurons). However, most LB23 interneurons showed overlapping connectivity with at least two BMN types innervating neighboring head bristle locations, while showing minimal LB23 connectivity overlap with BMNs from distant locations. In contrast, the LB23 descending neurons were less selective, receiving BMN inputs from broader head regions. This indicates that BMNs form somatotopically organized connections with LB23 neurons, ranging in resolution from specific bristle populations (e.g., proboscis) to broader head regions (e.g., ventral, dorsal, and posterior head, **Figure 14A,B**), supporting the existence of parallel BMN–LB23 pathways that correspond to head location.

**Figure 14.**
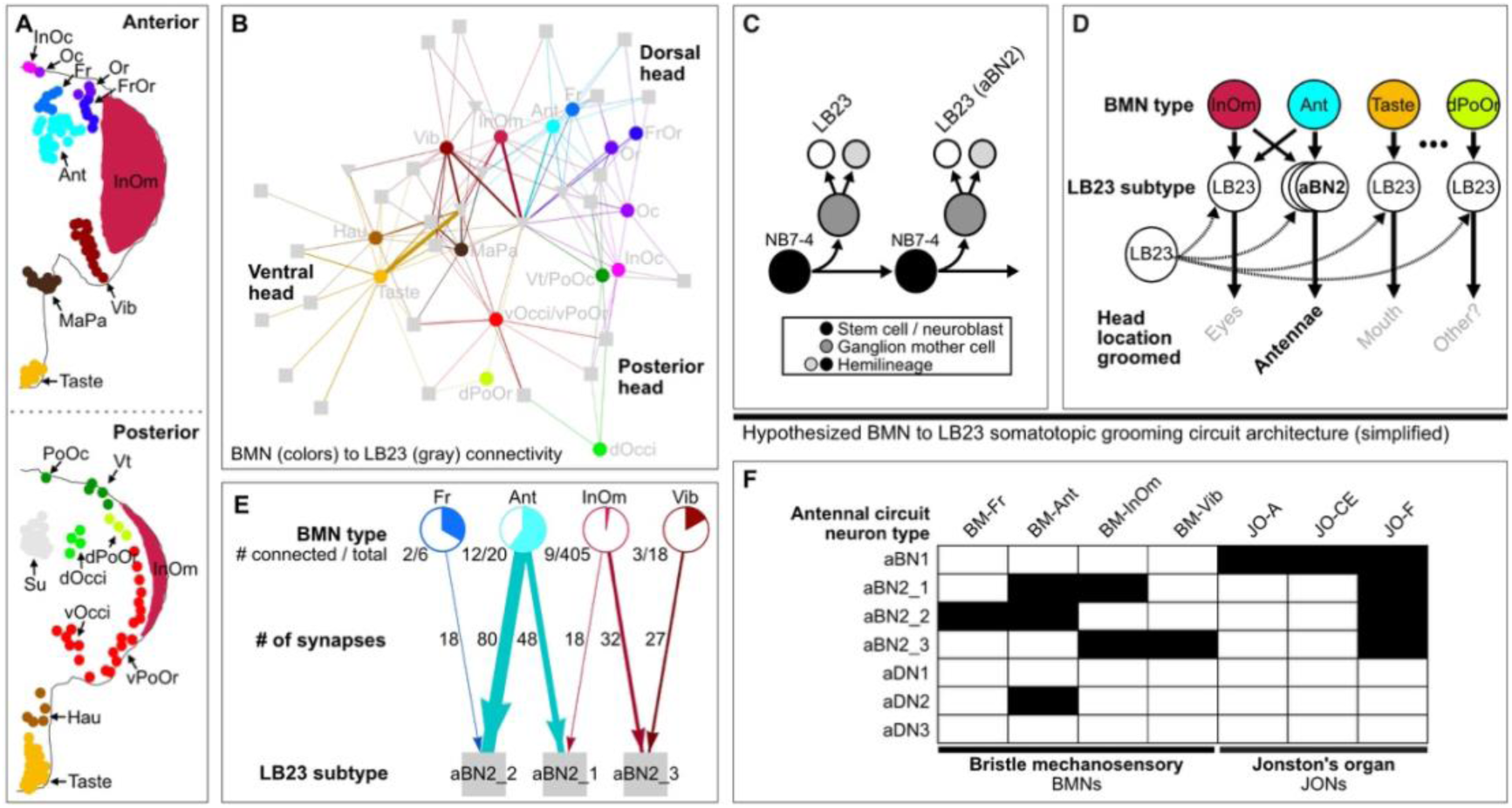
Developmentally related LB23 neurons form a putative somatotopically organized BMN architecture that underlies grooming. (**A**) Bristles on the anterior and posterior head, marked with labeled and color-coded dots indicating their classification. (**B**) Connections between BMN types (colored dots; synapses pooled) and their postsynaptic LB23 neurons (gray). Squares correspond to individual LB23 interneurons while triangles indicate descending neurons. BMN edges are weighted by synapse count and shown in their corresponding bristle colors in **A**. Bold text indicates the general head location of BMNs on the plot, revealing somatotopy-based connectivity with LB23 neurons (i.e. ventral, dorsal, and posterior head). Node positions were computed using a Fruchterman–Reingold force-directed layout: each directed edge acts like a spring whose strength is proportional to the edge weight (synapse count), drawing highly connected nodes together, while all nodes repel one another to prevent overlap. Consequently, neurons with many or strong synaptic connections cluster closely in the 2D embedding, whereas weakly or unconnected neurons are pushed to the periphery. (**C**) LB23 neurons are derived from NB7-4 neuroblasts that divide to produce another neuroblast and a ganglion mother cell. The ganglion mother cell divides once to produce an LB23 neuron and another neuron undergoing programmed cell death. Multiple rounds of this process produce the LB23 hemilineage that includes aBN2 neurons. (**D**) Hypothesized somatotopic BMN to LB23 parallel grooming circuit architecture. Each BMN type shows preferential connectivity with specific subpopulations of LB23 neurons. BMNs innervating neighboring bristles show overlapping connections with LB23 subpopulations. LB23 subpopulations elicit grooming of their corresponding presynaptic BMN locations. For example, aBN2 neurons are preferentially connected with BM-Ant neurons and elicit grooming of the antennae. (**E**) Connections of BMN types with antennal grooming LB23 neurons (previously named aBN2_1, _2, and _3). Shown connections have at least two neurons of a BMN type connected to the same aBN2 neuron, or are connected by greater than five synapses. Number of connected BMNs versus the total for each type is indicated and represented as a partially filled circle. Synapse numbers for each connection are indicated and represented by the width of the edges. **Supplementary file 16** provides a list of Flywire IDs for LB23 neurons which can be used to filter data from **Supplementary file 10**. (**F**) BMNs and JON types that are connected with neurons in the previously described antennal command circuit (black boxes). The BMN connections are reported in this work from Codex, whereas Johnston’s organ neuron (JON) connections were described previously (Hampel et al., 2020) (see Discussion).

Based on the results described above, we hypothesized that particular BMN head bristle locations are groomed with activation of subpopulations of postsynaptic LB23 hemilineage neurons (**Figure 14C,D**). If this is correct, then antennal BM-Ant neurons should be presynaptic to the LB23 subpopulation of aBN2 neurons that elicit antennal grooming (Hampel et al., 2015). Indeed, the majority of BMN output synapses onto aBN2 neurons were from BM-Ant neurons (**Figure 14E,F**), demonstrating that the somatotopic connectivity of BMN and aBN2 neurons corresponds with behavioral output. Further, BMN types innervating bristles from neighboring locations were also connected to aBN2 neurons, including BM-Fr, -InOm, and -Vib neurons. This suggests that neighboring BMN types could elicit antennal grooming through their connections with aBN2 neurons (**Figure 14D**).

While BM-Ant neurons showed the highest aBN2 connectivity among the BMN presynaptic partners, they only connected with two of the three aBN2 neurons, including aBN2_1 and _2, while aBN2_3 was connected with BM-InOm and -Vib neurons. This may indicate that aBN2_1 and _2 are connected mainly with BM-Ant neurons and elicit grooming of the antennae, while aBN2_3 has a role that is more specific for the BM-InOm and -Vib bristle locations. When taken together, our data uncovered a putative somatotopically organized and developmentally related circuit architecture that controls aimed grooming of the head (**Figure 14C,D**, see Discussion). This provides the foundation for studies that will define how spatially organized sensory input is transformed into coordinated motor output through lineage-derived neural pathways.

## Discussion

### Comprehensive BMN connectome organizational features reveal mechanosensory principles

A major advance of this work is the synaptic-resolution second-order connectome of *Drosophila* head BMNs, providing a comprehensive anatomical map of how peripheral touch signals interface with central circuits. Prior work established the functional and anatomical scaffolding for this map: activating BMNs from distinct head bristle populations evokes grooming of the corresponding head location, and nearly all head BMNs were assigned to their bristle populations and shown to project via distinct nerves into somatotopically organized zones in the ventral brain (Eichler et al., 2024). That study also provided initial evidence that BMNs from neighboring locations exhibit greater connectivity similarity to postsynaptic partners than BMNs from distant locations. What remained unresolved was a comprehensive description of BMN pre- and postsynaptic partners—including their anatomical diversity and the circuit-level principles underlying grooming behavior.

Here, we analyzed the reconstructed BMN connectome in the FAFB dataset (Dorkenwald et al., 2024; Eichler et al., 2024), focusing on second-order partners with ≥5 synapses onto or from BMNs. Even without incorporating fine-grained spatial resolution among BMN populations, this connectome reveals several dominant organizational features (**Figure 15A-C**).

**Figure 15.**
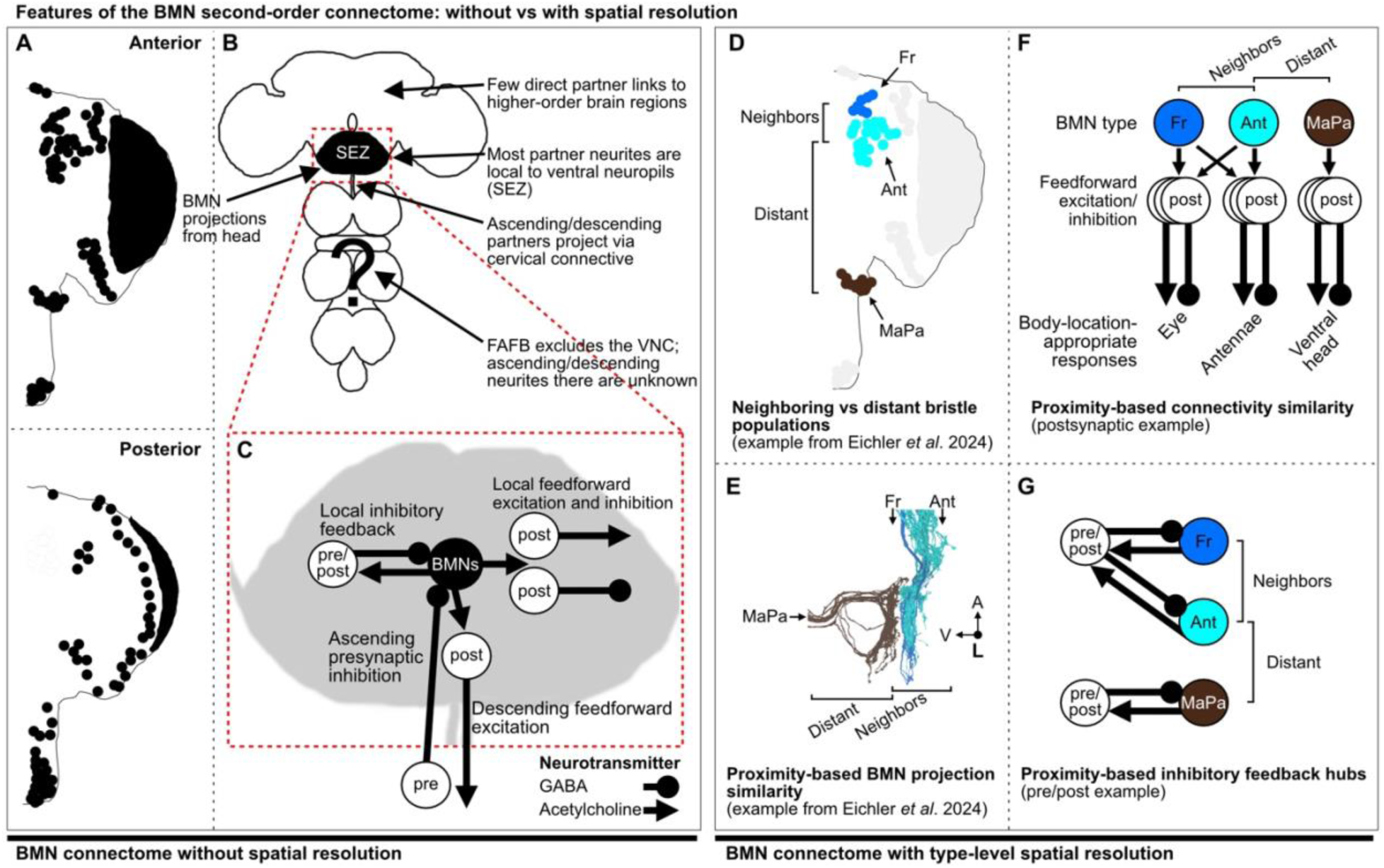
Features of the BMN second-order connectome: without vs with spatial resolution. (**A**) General features of mechanosensory processing were identified by analyzing the connections of nearly all BMNs innervating anterior and posterior head bristles (black dots), independent of spatial resolution of individual BMN populations. (**B**) Brain-level organization. BMNs project to the ventral brain neuropils, the gnathal ganglia (GNG; not shown). BMN partners rarely connect directly to dorsal higher-order regions because most partner neurites reside within the GNG and other ventral neuropils collectively referred to as the subesophageal zone (SEZ). Some partners have ascending or descending projections to/from the ventral nerve cord (VNC); however, the FAFB dataset does not include the VNC, so neurites within the VNC are not reconstructed. (**C**) Major SEZ-localized features (red box in **A**): local postsynaptic feedforward excitation and inhibition, descending postsynaptic feedforward excitation, ascending presynaptic inhibition, and local feedback inhibition. Partners are labeled “post,” “pre,” or “pre/post.” GABAergic neurons are shown as ball-and-stick symbols and cholinergic neurons as arrows. Features shown were prioritized by BMN pre- and postsynaptic synapse counts (largest fractions; Figures 7-8). (**D-G**) Spatially linking BMNs to their bristle populations reveals additional organizational features. (**D**) Head bristle/BMN populations include both neighboring and distant groups (Eichler et al., 2024). (**E**) BMNs project to somatotopically organized zones in the GNG, with neighboring populations overlapping more than distant populations (e.g., BM-Ant/BM-Fr neurons overlap; BM-MaPa neurons do not) (Eichler et al., 2024). (**F**) Using this spatial framework, the present study shows that BMN connectivity similarity is proximity-dependent, revealing somatotopically organized pre-and postsynaptic pathways. For example, neighboring BM-Ant and BM-Fr neurons share overlapping sets of connected postsynaptic partners, consistent with spatially organized excitatory and inhibitory pathways. Panel **F** was adapted and updated from Eichler et al. (2024). (**G**) The present study further identifies proximity-based inhibitory feedback hubs: neighboring BMNs show reciprocal connectivity with the same pre/post partners, indicating head-region-specific feedback inhibition (Figure 11). Panels **D** and **E** are reproduced from Eichler et al. (2024) (Figure 8D-F) under a CC BY license.

First, BMN circuitry is overwhelmingly ventral: BMNs project into the gnathal ganglia (GNG), and most pre-and postsynaptic partners concentrate their neurites in the GNG and neighboring subesophageal zone (SEZ) neuropils below the esophagus, with relatively sparse connectivity to dorsal higher-order brain regions (**Figure 15B**). This architecture suggests that early mechanosensory processing is largely confined to SEZ circuits, consistent with mechanosensory organization in other systems, including low-threshold mechanoreceptor circuits in the mammalian dorsal horn and mechanoreceptor pathways in locusts (Abraira et al., 2017; Newland and Burrows, 1997).

Second, inhibitory control is a defining feature of synaptic inputs onto BMNs. Presynaptic partners (excluding other BMNs) are predominantly predicted to be GABAergic, indicating that presynaptic inhibition is a major motif for scaling BMN output and sharpening spatial and temporal tuning (**Figure 15C**; (Blagburn and Sattelle, 1987; F. Clarac and Cattaert, 1996)). Many of these interneurons are also pre/post partners, embedding reciprocal loops within the afferent pathway and creating substrates for local feedback inhibition. A substantial fraction of presynaptic inhibitory inputs arise from neurons with ascending projections toward the brain, suggesting that body-state signals originating in the VNC gate head mechanosensation from below.

Third, BMN outputs engage diverse postsynaptic targets that can support both local processing and rapid behavioral control (**Figure 15C**). Postsynaptic partners are primarily predicted to be cholinergic, with additional GABAergic and glutamatergic populations, consistent with the heterogeneity of second-order mechanosensory partners described in vertebrate systems (Abraira et al., 2017). The connectome highlights prominent postsynaptic feedforward pathways within the SEZ, including excitatory and inhibitory branches, as well as strong engagement of descending neurons that receive a large fraction of BMN synapses and project to the VNC. This provides a direct anatomical route for head-derived touch signals to rapidly influence whole-body posture and reflexive actions. For example, stimulation of head BMNs can elicit rapid avoidance-like responses (Eichler et al., 2024), which could be mediated by descending neurons.

Together, these circuit features define an architectural framework that constrains early mechanosensory processing and links circuit organization to behavior. The ventral concentration of second-order partners confines early touch processing to SEZ circuits, limiting direct routing to dorsal regions. Within this framework, presynaptic inhibition provides a mechanism for transient suppression during hierarchical grooming sequences (Seeds et al., 2014), while recurrent pre/post loops offer candidate substrates for habituation and temporal integration of repeated touch (Corfas and Dudai, 1989; Guzulaitis et al., 2013; Sherrington, 1906; Stein, 2005). In parallel, descending projections provide a pathway through which head mechanosensory signals recruit whole-body motor programs.

By resolving BMN circuitry at synaptic resolution, this connectome links anatomical organization to specific circuit mechanisms and converts previously described behavioral phenomena into testable hypotheses. More broadly, it establishes the *Drosophila* head BMN system as a model for how peripheral touch signals are processed within localized hubs, regulated by inhibition, and distributed across parallel local and descending pathways to shape adaptive motor behavior.

Several limitations define priorities for future investigation. First, partners connected by fewer than five synapses were excluded, omitting weaker but potentially functional connections. For example, although 555 eye BM-InOm neurons are present in the FAFB left hemisphere, only 405 meet the five-synapse threshold (**Figure 2 – figure supplement 1B**). While individual BM-InOm neurons form relatively few synapses, their large numbers suggest they may collectively influence postsynaptic targets through spatial summation.

Second, the analysis focuses only on BMNs from the left hemisphere. Although contralateral neurons synapsing with left-side BMNs are included, the absence of the right-side BMN connectome limits assessment of bilateral symmetry, interhemispheric coordination, and side-to-side variability. Finally, because the FAFB dataset includes only the brain and not the ventral nerve cord (VNC), we cannot reconstruct VNC arbors of ascending or descending partners or identify their synaptic partners within the VNC (**Figure 15B**).

Forthcoming datasets spanning both brain and VNC will be required to address these gaps and define sensorimotor communication across the nervous system (Bates et al., 2025). Functional validation of the identified motifs will require calcium imaging, electrophysiology, and targeted perturbations. Nevertheless, by defining the pre-, post-, and reciprocal partners of a peripheral mechanosensory population at synaptic resolution, the BMN connectome provides a foundational framework for integrating anatomy, physiology, and behavior across the sensorimotor system.

### A synaptic resolution connectome of a head somatotopic map

Another striking feature is the extent to which the BMN connectome shows somatotopy with respect to different head locations. Prior work revealed that BMNs from the same bristle populations (same types) project into the same zones in the ventral brain, show high morphological similarity, and their morphology is stereotyped across individual flies (Eichler et al., 2024). Further, BMNs from neighboring bristle populations have overlapping projections into the same zones (**Figure 15D,E**). Here, we extend these findings by showing that this somatotopic organization is preserved at the level of synaptic distributions, further reinforcing the spatial structure of the head mechanosensory map (**Figure 1A-D**).

BMNs from different head locations are differentially represented by their synapse numbers in the brain (**Figure 1E**, **Figure 2A-C**). For example, BM-Taste neurons on the proboscis contribute the highest numbers of synapses among head BMN types, whereas BMNs on the posterior head contribute comparatively fewer synapses. BM-Taste neurons are distinct from other head BMNs in that many of their bristles are bichoid, containing both mechanosensory and gustatory neurons (Nayak and Singh, 1983; Shanbhag et al., 2001).

Consistent with this, BM-Taste neurons have been implicated in feeding and texture evaluation (Jeong et al., 2016; Sánchez-Alcañiz et al., 2017; Zhou et al., 2019), in addition to eliciting proboscis and ventral head grooming (Eichler et al., 2024). These multifunctional roles may account for their elevated synapse numbers. In contrast, BMNs innervating posterior head bristles may have more specialized functions, potentially explaining their lower synapse counts. The functional significance of these differences remains to be determined.

Importantly, somatotopy extends beyond BMN projections to their second-order synaptic partners. BMNs of the same type show the highest connectivity similarity, with progressively lower similarity observed among BMNs innervating neighboring versus distant head locations (**Figure 15F**). This organization is present in both pre- and postsynaptic partners (**Figure 10A-E**), as well as within the subset of pre/post partners alone (**Figure 11A**). Somatotopy therefore operates at multiple spatial scales, from individual bristle populations to broader head regions, including dorsal, ventral, and posterior regions. Overlapping connectivity among neighboring BMN type partners may enable responses to mechanosensory stimulations of neighboring bristles whose corresponding BMNs are likely to be stimulated together (Tuthill and Wilson, 2016b).

One notable exception to this pattern is the BM-InOm population, which occupies a central position in network diagrams and exhibits broad connectivity similarity with BMNs from across the head (**Figure 9A**, **Figure 10A-E**). This likely reflects the large surface area of the compound eyes, which span dorsal, ventral, and posterior regions and neighbor multiple bristle populations. Consistent with previous work showing morphological diversity among BM-InOm neurons (Eichler et al., 2024), our output connectivity analysis suggests the presence of multiple BM-InOm subtypes defined by distinct partner profiles (**Figure 9A**). Future work will be needed to determine how this heterogeneity relates to spatial organization within the eye.

Somatotopic connectivity was observed across diverse BMN partners, including putative primary neurons, which function in both larval and adult stages, and secondary hemilineages, which are specific to the adult brain (**Figure 12 – figure supplement 1A-G**). In this study, we focused on two partner populations: the cholinergic hemilineage LB23 and a set of bilaterally symmetric GABAergic pre/post neuron pairs involved in BMN feedback inhibition (**Figure 15G**). Both populations exhibited connectivity patterns that mirrored the broader somatotopic organization of the BMN connectome, with preferential coupling to BMNs from specific head locations (**Figure 10A-E**, **Figure 11B-I**, **Figure 14A,B**). Whether somatotopy is a general property of all BMN second-order partners remains an open question for future investigation.

### First comprehensive connectome of somatotopically-organized mechanosensory neurons

Somatotopic maps have been identified across species, as exemplified by vertebrate head and body maps (Abraira and Ginty, 2013; Adibi, 2019; Brown et al., 1977). Somatotopic organization is preserved through different processing layers in the nervous system and thought to be fundamentally important, although the functional significance of this organization remains unknown (Kaas, 1997; Thivierge and Marcus, 2007). High-resolution anatomical maps are therefore essential for defining how somatotopic organization is implemented at the circuit level and how it influences behavior.

Previous studies in both vertebrates and invertebrates have examined aspects of mechanosensory somatotopy, including receptive field organization, second-order neuron diversity, and behavioral relevance (Abraira et al., 2017; Abraira and Ginty, 2013; Brown et al., 1987, 1980, 1977; Newland and Burrows, 1997). These studies revealed striking parallels in the integration of mechanosensory inputs across phyla, but were necessarily limited in scope by the lack of complete synaptic-resolution datasets. As a result, comprehensive descriptions of second-order mechanosensory circuit architectures have remained elusive.

By leveraging a full adult brain connectome, this work provides a synaptic-resolution view of head somatotopic mechanosensory circuitry with previously defined behavioral outputs (Eichler et al., 2024). The BMN connectome described here reveals fundamental organizational principles—including parallel somatotopic pathways, region-specific excitatory and inhibitory outputs, and extensive inhibitory control—that are likely to generalize across mechanosensory systems (**Figure 15D-G**). This dataset establishes a foundation for future studies aimed at linking somatotopic circuit architecture to sensorimotor transformations and action selection, and for testing how parallel sensory pathways contribute to flexible, context-dependent behavior.

### Hypothesis: LB23 neurons translate the head BMN somatotopic map into aimed grooming movements

Flies can groom specific locations on their head, such as the eyes, antennae, proboscis, dorsal and ventral head (Dawkins and Dawkins, 1976; Hampel et al., 2017, 2015; Seeds et al., 2014; Szebenyi, 1969; Zhang et al., 2020). These movements can be elicited by stimulating specific head bristles or through optogenetic activation of specific BMN types (Eichler et al., 2024; Hampel et al., 2017; Melzig et al., 1996; Zhang et al., 2020). However, the range of aimed movements that are elicited by stimulating each bristle population remains to be experimentally determined. Head BMNs project from bristles to somatotopically organized zones in the ventral brain, with those innervating neighboring bristle populations occupying overlapping zones (**Figure 1A-D**). To understand how the BMNs elicit aimed head grooming, we examined their postsynaptic targets. We identified hemilineage 23b (LB23) neurons as important mediators of BMN-driven, aimed head grooming.

LB23 neurons were previously suggested to be connected with BMNs based on their anatomical overlap (Harris et al., 2015; Shepherd et al., 2019), and here we provide direct evidence of this connectivity. LB23 is the only hemilineage in which 100% of its members are connected with ipsilateral BMNs, suggesting a critical role in the processing of BMN inputs (**Figure 12A,B**). The BMN to LB23 connectivity network is somatotopic, showing overlapping connectivity among neighboring head BMNs, however, not among distant ones (**Figure 14B**). Thus, the BMNs and LB23 neurons form different parallel pathways that correspond to head location.

In this work, we identified the LB23 hemilineage members among the neuron types previously shown to form a neural circuit that elicits grooming of the antennae in response to mechanical stimulations (Hampel et al., 2020b, 2015). The circuit receives inputs from mechanosensory chordotonal organ neurons called Johnston’s organ neurons (JONs) that detect mechanical displacements of the antennae and project to the ventral brain. There, they excite two brain interneuron types, including one aBN1 neuron and three aBN2 neurons, along with three descending neurons (aDNs). aDNs project to the VNC where they are hypothesized to engage pattern generation circuitry that produces antennal grooming leg movements (Berkowitz and Laurent, 1996).

aBN2 was the only type in the antennal grooming circuit with morphological similarity to LB23 neurons. We used the MCFO technique to label individual aBN2s, showing that their morphologies were preserved across individuals (**Figure 13D-F**). Two aBN2s were indistinguishable from each other, while the third had a midline projecting branch. The reconstructed LB23 neurons consist of different morphologically distinct neurons, and three of these neurons showed striking similarity to the individually labeled aBN2s, including one with a midline projecting branch (**Figure 13D-I**). This linked previously studied aBN2s to the LB23 hemilineage. As further confirmation that the three reconstructed LB23 neurons are aBN2, they are all postsynaptic to aBN1 in FAFB, and thus a part of the antennal grooming circuit (Hampel et al., 2020b, 2015). This identifies a subpopulation of three morphologically similar LB23 neurons as aBN2s that elicit aimed grooming of the antennae.

According to our model of grooming (**Figure 1 – figure supplement 1A,C**), BMN/LB23 connectivity should form parallel pathways that elicit grooming corresponding to BMN location. For example, the aBN2s that elicit antennal grooming should be postsynaptic to antennal BMNs. Indeed, antennal BM-Ant neurons are the most highly aBN2-connected BMNs with two aBN2 neurons, aBN2_1 and _2 (**Figure 14E,F**). Thus, the location groomed (i.e. antennae) corresponds to the location of the majority of BMN inputs. However, aBN2_1 and _2 also have lower level connections with BMN types from neighboring locations, including the BM-Fr, -InOm neurons. This overlap could explain our previous finding that optogenetic activation of specific BMN types elicits aimed grooming of their corresponding head bristle locations, but also neighboring locations (Eichler et al., 2024). An organization whereby neighboring BMNs connect with aBN2 neurons that elicit grooming of neighboring locations provides a means of cleaning locations neighboring the stimulus site. One caveat stems from the finding that BM-Ant neurons do not connect with aBN2_3 that is instead connected with BM-ImOn and -Vib neurons from locations neighboring the antennae. This raises the possibility that activation of aBN2_3 elicits grooming of areas neighboring the antennae, such as a ventral location where the BM-InOm and -Vib neurons are located, which might be impossible to distinguish by eye (**Figure 14A**). Testing this will require high resolution behavioral analysis of the grooming elicited by activating individual aBN2 neurons.

While our past and present work together reveal that a subpopulation of LB23 neurons elicits antennal grooming, we also find evidence that other LB23 neurons in the hemilineage elicit additional head grooming movements. We previously reported a driver line (R40F04-GAL4) that elicits grooming of the whole head (Seeds et al., 2014). Here, we show that this line expresses in a cluster of 14 ventral brain neurons whose collective morphology resembles that of the LB23 neurons (**Figure 13B,C**). While previous work did not distinguish between the LB23 neurons and other neurons targeted by R40F04-GAL4 as the ones eliciting grooming (**Figure 13B**), our unpublished work implicates LB23 as the relevant neurons in this pattern. Thus, collective activation of at least 14 LB23 neurons elicits grooming of the whole head, implicating LB23 hemilineage neurons, in addition to the previously described aBN2s, as drivers of head grooming.

We hypothesize that the BMNs and LB23 hemilineage neurons form a somatotopic parallel grooming circuit architecture that elicits grooming of different head locations in response to mechanical stimulations (**Figure 14D**). LB23 neurons are morphologically diverse, and a preliminary assessment indicates that these neurons could be categorized into subpopulations based on morphological similarity (**<u>FlyWire.ai link 1</u>**), as we found with the aBN2 subpopulation (**Figure 13D-I**). The different BMN types that are preferentially connected with subpopulations of LB23 neurons are hypothesized to elicit aimed grooming of their corresponding bristle locations. Further, BMNs innervating neighboring bristles show overlapping connections with LB23 subpopulations, providing a mechanism by which bristle stimulation results in grooming of the stimulus location, and neighboring locations. Future experiments will define the full morphological diversity of the LB23 neurons and test the extent to which activation of LB23 subpopulations elicits aimed grooming of different head locations.

This work provides evidence of a somatotopically organized parallel circuit architecture that is established by a developmentally related set of neurons. Further, at least one subpopulation of LB23 neurons (aBN2 neurons) elicits grooming of the location of the BMN that they are connected to (BM-Ant neurons) (**Figure 14E,F**), and we hypothesize that other LB23 neurons also elicit different aimed head grooming movements (**Figure 14D**). This supports a model where LB23 subpopulations are developmentally pre-committed to body-location-specific sensorimotor roles. In addition to being in the ventral brain, LB23 neurons are also in different segments in the VNC, where they are thought to be connected with BMNs from different body locations (Harris et al., 2015; Shepherd et al., 2019). Previous studies showed that stimulations to bristles on the body also elicit aimed grooming locations including the legs, wings, and thorax (Corfas and Dudai, 1989; Li et al., 2016; Matheson, 1997; Page and Matheson, 2004; Usui-Ishihara et al., 1995; Vandervorst and Ghysen, 1980). Future studies will be necessary to test the BMN/LB23 connectivity and functional organization in both the ventral brain and VNC.

### The antennal grooming circuit receives diverse mechanosensory inputs

Our previous work and the present study reveal that the antennal grooming circuit receives inputs from two different classes of antennal mechanosensory neurons, the BMNs and JONs. We previously reported that different subpopulations of JONs converge onto the antennal grooming circuit and elicit grooming (Hampel et al., 2020b, 2020a, 2015). These subpopulations are morphologically distinct, and respond to different types of antennal stimulations. They include JO-A neurons that respond to high frequency vibrations, JO-CE neurons that respond to antennal displacements and vibrations, and JO-F neurons that do not respond to any of these stimuli, whose activation stimulus remains unknown (Hampel et al., 2020a; Kamikouchi et al., 2009; Mamiya and Dickinson, 2015; Matsuo et al., 2014; Patella and Wilson, 2018; Yorozu et al., 2009). While JO-A neurons have not been tested, optogenetic activation of either JO-CE or JO-F neurons elicits antennal grooming. The different JON subpopulations, aBN1, aBN2s, and aDNs neurons were previously identified in the FAFB dataset (Hampel et al., 2020b; Shiu et al., 2024). This revealed that JO-A, -CE, and -F neurons are all connected with aBN1, while only JO-F neurons are connected with aBN2 and aDN2 (**Figure 14F**). Here, we find that the BMNs are also connected with the antennal circuit, implicating another mechanosensory modality involved in eliciting grooming behavior. The aBN2 neurons were shown to receive most BMN inputs in the antennal grooming circuit, although aDN2 also had connections with BMNs (**Figure 14F**). Thus, different types of mechanical stimuli may elicit antennal grooming.

### Parallel circuit architecture underlying the grooming sequence

The mechanosensory neurons hypothesized from the parallel model that elicit the *Drosophila* grooming sequence were identified in previous work (Eichler et al., 2024; Hampel et al., 2020a, 2017, 2015; Mueller et al., 2019; Seeds et al., 2014; Zhang et al., 2020). These neurons are at different locations on the body and head and elicit aimed grooming of their corresponding bristles. The different movements are performed in a sequence when mechanosensory neurons at different locations become simultaneously activated by dust covering the body. Supporting this, simultaneous optogenetic stimulation of mechanosensory neurons at these locations induces a sequence resembling natural dust-induced grooming (Hampel et al., 2017; Zhang et al., 2020). BMNs play a pivotal role, as their activation alone can initiate aimed grooming, whereas their collective activation elicits a grooming sequence (Zhang et al., 2020).

Here, we focus on BMN postsynaptic partners that could form parallel circuits that elicit aimed grooming (**Figure 1 – figure supplement 1A,C**). Consistent with a parallel architecture, we identified subpopulations of LB23 neurons showing preferential connectivity with BMN types innervating neighboring head bristles (**Figure 14A,B**). At least some LB23 neurons elicit aimed head grooming, as we demonstrated by linking previous optogenetic activation experiments with the LB23 anatomical comparisons (**Figure 13A-I**). This leads us to hypothesize that different BMN types connect with morphologically distinct LB23 subpopulations, establishing a parallel architecture that elicits aimed head grooming (**Figure 14D**).

The spatial resolution of the sensory-to-motor transformation in this parallel circuit architecture remains to be tested. While grooming is aimed at specific head locations in response to mechanical stimulations, grooming of neighboring locations can also be elicited (Eichler et al., 2024; Hampel et al., 2017, 2015). The extent to which different BMN types elicit grooming of distinct locations versus neighboring locations remains to be systematically defined. The BMN to LB23 connectivity may offer insights into the circuit architecture underlying these responses, as neighboring types have overlapping connectivity with LB23 neurons (**Figure 14A,B**). Thus, stimulation of a particular BMN type could excite multiple postsynaptic neurons, potentially eliciting grooming of local and neighboring locations. Future studies will test this by linking the connectivity with high resolution analysis of the grooming movements elicited with activation of different nodes in the BMN circuit architecture.

Simultaneous activation of the parallel circuit architecture by dust causes competition among the movements to be performed in sequence (Seeds et al., 2014). The competition is resolved through hierarchical suppression, where earlier movements in the sequence suppress later movements (**Figure 1 – figure supplement 1B**). An activity gradient is established among the parallel circuits to determine performance order where earlier movements have the highest activity and later ones have the lowest. The highest activity movement is selected by a winner-take-all network that suppresses the other movements. The activity gradient could be established through the control of sensory gain among the BMNs, or through lateral inhibitory connections between the parallel circuits. We previously hypothesized that this suppression could occur through unilateral inhibitory connections between the different parallel circuits or by modulating the sensory gain on the different grooming circuits (Seeds et al., 2014).

Work presented here reveals one possible mechanism of sensory gain control through presynaptic inhibition of the BMNs. Indeed this mechanism has been shown to control different movements by negatively regulating mechanosensory output (Blagburn and Sattelle, 1987; Burrows and Matheson, 1994; F Clarac and Cattaert, 1996; Gaudry and Kristan, 2009). Here we find that most of the presynaptic inputs onto the BMNs are GABAergic, further implicating presynaptic inhibition in controlling grooming behavior. These GABAergic neurons are somatotopically organized, showing preferential connectivity with BMNs from specific head locations (**Figure 10B,C**, **Figure 11A**). Somatotopy-based presynaptic inhibition could establish the activity gradient across the different BMN parallel pathways described above. The presynaptic inhibitory neurons identified here enable future studies to test the roles of these neurons in mediating presynaptic inhibition and controlling grooming behavior.

## Methods

### Key resources table

| Reagent type (species) or resource | Designation | Source or reference | Identifiers | Additional information |
| --- | --- | --- | --- | --- |
| Data platform | Codex | Matsliah *et al.* 2003 |  | http://dx.doi.org/10.13140/RG.2.2.35928.67844 |
| Data platform | Flywire-Codex | Dorkenwald *et al.*, 2024 | Flywire v783 | https://codex.flywire.ai |
| Software, algorithm | R | R Core Team | v4.2.2 | https://www.r-project.org/ |
| Software, algorithm | natverse | Bates *et al.*, 2020 | v0.2.4 | http://natverse.org/ |
| Software, algorithm | fafbseg | Bates *et al.*, 2020 | v0.14.0.90 | Part of natverse |
| Software, algorithm | igraph | Csardi and Nepusz, 2006 | v2.1.4 | https://igraph.org/r/ |
| Software, algorithm | ggplot2 | Wickham, 2016 | v3.5.1 | https://ggplot2.tidyverse.org/ |
| Software, algorithm | visNetwork | Almende B.V. & Contributors | v2.1.3 | https://datastorm-open.github.io/visNetwork/ |
| Software, algorithm | networkD3 | Allaire et al. | v0.4 | https://christophergandrud.github.io/networkD3/ |
| Software, algorithm | sunburstR | Russell et al. | v2.1.8 | https://github.com/timelyportfolio/sunburstR |
| Software, algorithm | heatmaply | Galili et al., 2018 | v1.5.0 | https://cran.r-project.org/package=heatmaply |
| Software, algorithm | Isa | Wild, 2022 | v0.73.3 | https://cran.rstudio.com/web/packages/Isa/ |
| Software, algorithm | Python | Python Software Foundation | v3.10.9 | https://www.python.org/ |
| Software, algorithm | navis | Schlegel et al., 2024 | v1.9.1 | https://navis-org.github.io/navis/ |
| Software, algorithm | Blender | Blender Foundation | v3.0.1 | https://www.blender.org |
| Software, algorithm | Tableau | Salesforce/Tableau Software | 2023.2.1 | https://www.tableau.com/ |
| Software | Fiji | Schindelin et al., 2012 |  | http://fiji.sc/ |
| Genetic reagent (D. melanogaster) | R404F04 - GAL4 | Seeds et al., 2014 |  |  |
| Genetic reagent (D. melanogaster) | aBN2-spGAL4-1 | Hampel et al., 2015 |  |  |
| Genetic reagent (D. melanogaster) | *10XUAS-IVS-mCD8::GFP* | Pfeiffer et al., 2010 | RRID: BDSC_32194 |  |
| Genetic reagent (D. melanogaster) | MCFO-5 | Nern et al., 2015 | RRID: BDSC_64089 |  |
| Antibody | anti-GFP (Rabbit polyclonal) | Thermo Fisher Scientific | Cat#A-11122, RRID:AB_221569 | IF (1:500) |
| Antibody | anti-Brp (Mouse monoclonal) | DSHB | Cat#nc82, RRID:AB2314866 | IF (1:50) |
| Antibody | anti-FLAG (Rat monoclonal) | Novus Biologicals | Cat#NBP1-06712, RRID:AB_1625981 | IF (1:300) |
| Antibody | anti-HA (Rabbit monoclonal) | Cell Signaling Technology | Cat#3724, RRID:AB_1549585 | IF (1:500) |
| Antibody | anti-V5<br>(Mouse monoclonal) | BIO-RAD | Cat#MCA1360,<br>RRID:AB_322378 | IF (1:300) |
| Antibody | anti-Rabbit<br>AF488<br>(Goat polyclonal) | Thermo<br>Fisher<br>Scientific | Cat#A-11034,<br>RRID:AB_2576217 | IF (1:500) |
| Antibody | anti-Mouse<br>AF568<br>(Goat polyclonal) | Thermo<br>Fisher<br>Scientific | Cat#A-11031,<br>RRID:AB_144696 | IF (1:500) |
| Antibody | anti-Rat<br>AF633<br>(Goat polyclonal) | Thermo<br>Fisher<br>Scientific | Cat#A_21094,<br>RRID:AD_2535749 | IF (1:500) |

### Connectome data and neuron meshes

Based on previous work (Eichler et al., 2024), bristle mechanosensory neurons (BMNs) were proofread, categorized, and compiled into a master list of left-sided BMNs (previously classified as right-sided BMNs) with their corresponding root ID in Flywire (v783) (Dorkenwald et al., 2024; Zheng et al., 2018) as seen in **Supplementary file 1**. Synaptic connectivity data for BMNs and all pre- and postsynaptic partners with five or greater connections were obtained from the FlyWire Codex platform (https://codex.flywire.ai/), which provides preprocessed reconstructions and synapse annotations across the *Drosophila* brain (Buhmann et al., 2021; Dorkenwald et al., 2024; Heinrich et al., 2018; Zheng et al., 2018).

Synapse counts throughout this study are based on FlyWire/Codex synapse annotations and represent the number of individual pre- to postsynaptic contacts (incoming or outgoing connections), rather than the number of presynaptic active sites (T-bars); thus, presynaptic counts reflect polyadic connectivity as described previously (Schlegel et al., 2024).

The full connections.csv data table was downloaded from Codex (https://codex.flywire.ai/). This table contains one row for each pair of connected neurons within a given neuropil and reports the number of synapses between those neurons. Only connections involving five or more synapses (combined across all neuropils) are included in this dataset; pairwise connections with fewer than five synapses are excluded (Dorkenwald et al., 2024). Therefore, connections between BMNs and any partners with fewer than five synapses were not included in any analysis present in this work.

To identify BMN connectivity, the table was filtered for entries in which either the presynaptic or postsynaptic root ID (v783) matched a left-sided BMN. Rows with BMNs as the presynaptic root ID were classified as BMN outputs, and the postsynaptic root ID was classified as a postsynaptic partner of BMNs (**Supplementary file 2**). Similarly, rows with BMNs as the postsynaptic root ID were classified as BMN inputs, and the presynaptic root ID was classified as a presynaptic partner (**Supplementary file 3**). Unique partner IDs were compiled from the filtered inputs and outputs (**Supplementary file 6**). Partners that appeared as both pre- and postsynaptic to BMNs were categorized as “pre/post neurons” enabling us to quantify reciprocal interactions and assess the proportion of BMN inputs that originated from their downstream targets.

Three-dimensional neuron meshes for BMNs and their partners were obtained from cloudvolume data in R (v4.2.2) using the fafbseg package and the broader natverse framework (Bates et al., 2020) (**Figure 1C,D**, **Figure 1 – figure supplement 2**). The brain surface mesh used for anatomical context was also imported from the fafbseg package in R. Synapse locations were obtained from the synapse_coordinates.csv table from Codex (Dorkenwald et al., 2024) and plotted alongside meshes to visualize the distribution of pre- and postsynaptic sites of BMNs (**Figure 1C,D**, **Figure 1 – figure supplement 3**). To maintain consistency with FlyWire.ai, all neurons in this manuscript are displayed in the opposite brain hemisphere (e.g left hemisphere neurons are shown on the right) (Dorkenwald et al., 2024).

### Quantification of BMN synaptic input and output

Synaptic input and output counts were quantified for each individual BMN and grouped by BMN subtype. Data were obtained from manually curated summary tables containing total presynaptic and postsynaptic synapse counts for each BMN (**Supplementary file 1**) and weighted connections between BMNs and their partners (**Supplementary file 2** and **Supplementary file 3**). The total number of inputs and outputs found for each BMN of a particular type were visualized using an area plot generated by importing the data into Tableau (**Figure 1E**). These values were also used to generate a visualization of BMN synapse distribution across the head (**Figure 2**), as well as a visualization of synapse counts for each individual BMN within a particular BMN type (**Figure 2 – figure supplement 1A,B**). Box plots in **Figure 2 - figure supplement 1A** were generated separately for inputs and outputs in R using the ggplot2 package (Wickham, 2016). A secondary axis was included to accommodate the dual representation of input and output counts on a shared x-axis.

### Categorization of BMN partners

We summarized the total number of synapses for each BMN type, broken down into inputs and outputs, and quantified the number of unique synaptic partners. To gain an overview of partner identity, all presynaptic and postsynaptic partners of BMNs – including partners classified as “pre/post” – were previously inspected manually in FlyWire and categorized into five broad morphological classes: interneurons, descending neurons, ascending neurons, other BMNs, and motor neurons. These classes were cross-referenced with existing annotations in FlyWire (Dorkenwald et al., 2024).

Neurons with both dendritic and axonal processes, as well as a soma, entirely within the CNS were categorized as interneurons (noted as “Central Brain” or “CB” in flywire annotations). Neurons with their soma and dendritic processes in the CNS and axonal projections extending into the descending tracts of the VNC were categorized as descending neurons. Conversely, neurons originating from the cervical connective that contained axonal projections in the CNS were classified as ascending neurons.

To further characterize BMN partners, the full neurons.csv table was downloaded from Codex. This table provides a row for each neuron in the dataset, including its neurotransmitter prediction (Buhmann et al., 2021; Dorkenwald et al., 2024; Heinrich et al., 2018; Zheng et al., 2018). Partner root IDs were cross-referenced against this table to assign a predicted neurotransmitter identity to each BMN partner (**Supplementary file 6)**.

### Visualization of BMN partner category and synapse composition

Partner categories and connectivity patterns with BMNs were visualized using a Sankey diagram (i.e., directional flow diagram) (**Figure 3A, Supplementary file 4**, **Supplementary file 5**) and two sunburst plots (**Figure 3B**, **Supplementary file 6**) generated in R. In **Figure 3A**, normalized synapse counts between each BMN type and its presynaptic or postsynaptic partners were used. To achieve this, BMN synapses were grouped by target partner class (e.g., interneurons, descending neurons, etc.) and further subdivided into purely presynaptic, purely postsynaptic, or pre/post categories (**Supplementary file 4)**. Synapse values were normalized within each BMN type by setting all edge weights proportional to the total number of outputs for the particular BMN type (**Supplementary file 5)**. This allowed for direct comparison of synapse fractions across BMN types. The normalized matrix was reshaped into a long-format edge list and used to construct a directional flow diagram using the networkD3 package in R (**Figure 3A**).

To further characterize the composition of BMN synaptic partners and their associated synapses, sunburst plots were created using the sunburstR package in R (**Figure 3B**). One plot represented the distribution of presynaptic, postsynaptic, and pre/post partners by category and neurotransmitter identity; the other depicted the same structure based on the number of pre- and postsynaptic sites rather than partner counts. Each sunburst plot consisted of three layers (Schlegel et al., 2021): the first denoting a pre- or postsynaptic (or pre/post) relationship with BMNs; the second indicating morphological class (e.g. interneuron, ascending); and the third showing predicted neurotransmitter identity (Dorkenwald et al., 2024; Eckstein et al., 2024; Matsliah et al., 2023). Neurons for which neurotransmitter predictions were not available were labeled as “unknown”.

To visualize the brain occupying space of different partner categories, three-dimensional meshes of all partners were downloaded using fafbseg in R and displayed alongside BMNs for anatomical reference (**Figure 5**). Here, partners were separated based on category (e.g., interneuron, ascending) as well as their relationship to BMNs (e.g., presynaptic, postsynaptic). Further analysis was performed to highlight the varying neurotransmitter profiles and brain occupying regions of presynaptic and postsynaptic partners (**Figure 7**, **Figure 8, Supplementary file 6**).

### Identification and visualization of BMN-connected motor neurons

Motor neurons postsynaptic to BMNs were identified using annotations available in the Codex platform. For each motor neuron, the number of presynaptic BMNs was quantified. Connections were summarized in a matrix representing the number of individual BMNs of each type that were presynaptic to each motor neuron (**Figure 4, Supplementary file 7**). This matrix was visualized as a heatmap, with grayscale intensity indicating the number of presynaptic BMNs per motor neuron. Neurons were grouped by the peripheral nerve through which their axons exit the brain: labial, pharyngeal, or antennal, as annotated in FlyWire. These categories were color-coded for both heatmap visualization and mesh rendering (**Figure 4C**).

Three-dimensional meshes of all postsynaptic motor neurons were downloaded using fafbseg in R and displayed alongside BMNs for anatomical reference (**Figure 4A**). Anterior, dorsal, and lateral views were rendered to show the spatial distribution of each motor neuron class within the brain (**Figure 4B**). To visualize the peripheral organization of BMNs projecting to each motor nerve class, colored dots were overlaid on bristle maps, marking the head regions innervated by BMN types presynaptic to each class (**Figure 4D-F**). In addition to raw BMN counts, a second heatmap was generated to compare the percent of total synaptic input each motor neuron received from each BMN type (**Figure 4 - figure supplement 1**).

### Synaptic distribution of partners across neuropils

Using the Codex neuropil_synapse_table.csv table (https://codex.flywire.ai/), we assessed the number of input and output synapses across brain neuropils for all BMN partners categorized as ascending, descending, or interneuron. This table includes a column with a neuron’s root ID and the number of synapses in each neuropil of the brain as a separate column. The root ID column of this table was filtered to include only root IDs matching the BMN partners (**Supplementary file 8**).

Partners were grouped into three categories: purely presynaptic, purely postsynaptic, and pre/post partners. For each partner category, synapses were grouped by neuropil based on the location annotation provided in the Codex dataset (**Supplementary file 9**). This enabled identification of the upstream input sources of BMN presynaptic partners and the downstream output targets of BMN postsynaptic partners. Partners with both pre- and postsynaptic connections to BMNs were analyzed separately, allowing us to assess synaptic distributions for purely presynaptic, purely postsynaptic, and bidirectionally connected partner populations independently.

Given the overwhelming concentration of synapses (82%) in the gnathal ganglia (GNG), data were normalized to emphasize distribution patterns outside of this region. Specifically, for each partner category, the neuropil with the second-highest synapse count was used as the normalization reference. This neuropil was assigned a value of 1.0, and synapse counts in all other neuropils were expressed as fractions of that value. By excluding the GNG from normalization and referencing the second-most prevalent neuropil, this approach avoids compression of signal in other regions due to GNG dominance and enables clearer comparison of neuropils outside the GNG.

For visualization of neuropil distributions, normalized synapse data were merged with neuropil surface meshes in Blender (**Figure 6**). Neuropil meshes and the full brain mesh were imported into Blender’s Python API using the navis package (Schlegel et al., 2024). A custom Python script applied color values to neuropil mesh materials based on the normalized synapse fractions using a linear interpolation between white (low) and red (high) values. Separate interpolation functions were applied to base color, emission, transmission, and transparency values to enhance visualization results. Neuropils with no associated synapses were automatically hidden from both the viewport and render output.

Normalized data used for visualization and raw synapse counts are available in **Supplementary File 9**. Neuropils and images were colored and rendered independently for each partner group (presynaptic, postsynaptic, and pre/post). For each panel in **Figure 6** color intensity reflects synapse density relative to the second-most innervated region in each respective group.

### Network visualizations

Network graphs were generated separately for BMN presynaptic and postsynaptic connections using Codex-derived data (**Figure 10B-E**). The filtered connectome data obtained from the Codex connections.csv table in the form of an edge list (**Supplementary file 10**) was converted into two bipartite adjacency matrices: one for BMNs and their presynaptic partners, and another for BMNs and their postsynaptic partners. Neurons classified as “pre/post” were included in both network graphs.

Each adjacency matrix was imported into R and converted into a directed weighted graph using the igraph package (Csárdi and Nepusz, 2006). Edge weights reflected the number of synapses between BMNs and their partners. Node metadata, including classification of BMNs and their partners, was obtained from manually curated annotation tables (**Supplementary file 1, Supplementary file 6**).

Network graphs were visualized using the visNetwork package and its interface with igraph in R. The visIgraphLayout function was used to generate spatial node arrangements based on the force-directed layout algorithm by Fruchterman-Reingold, with a fixed random seed (1234) to ensure consistency across plots.

Node shapes were assigned based on partner categories (BMNs, interneuron, descending, or ascending), and edge thickness was scaled to synapse count. Node positions were determined by the Fruchterman–Reingold force-directed layout, in which each directed edge acts like a spring whose strength is proportional to the edge weight, drawing highly connected nodes together, while all nodes repel one another to prevent overlap. Consequently, neurons with many or strong synaptic connections cluster closely in the 2D embedding, whereas weakly or un-connected neurons are pushed to the periphery.

The relative spatial positioning of nodes in the network layouts was used to evaluate the organization of BMN types based on their corresponding anatomical origins on the head. Nodes corresponding to individual BMN types were grouped according to their associated head bristle locations as belonging to the dorsal, ventral, or posterior portion of the head, and their positions in the network were assessed to determine whether these groups exhibited spatial separation or clustering (**Figure 10C,E**).

### Cosine similarity clustering of individual BMNs

To assess the similarity of BMNs based on their synaptic outputs, pairwise cosine similarity scores were calculated using an adjacency matrix constructed from Codex synapse data. The matrix was derived from a directed edge list (**Supplementary file 11**) in which each row represented the presence of a connection from a BMN to a postsynaptic partner. The adjacency matrix was binarized such that entries indicated the presence (1) or absence (0) of a synaptic connection, without incorporating synaptic weight. The matrix was transposed so that each BMN was represented as a vector of its output connections, and cosine similarity was computed between all BMN vectors using the lsa package in R (**Supplementary file 12**).

To reduce background and emphasize stronger pairwise relationships, we applied a threshold of 0.3 to the similarity scores, with all values below this threshold set to zero. The resulting matrix was visualized as a clustered heatmap using the heatmaply package (Galili et al., 2017) with Ward’s hierarchical clustering method (**Figure 9A**). Color bars were included to indicate BMN subtype and corresponding anatomical head position.

BMNs classified as interommatidial (InOm) were excluded from a second round of plotting to better visualize the relationships between other BMN subtypes. The previously computed similarity matrix was subset by removing rows and columns corresponding to InOm BMNs, and a second heatmap was generated including all the remaining BMN types (**Figure 9B)**.

The thresholded similarity matrix was also used to construct an undirected weighted network graph using the igraph package in R (**Figure 9E)**. A force-directed layout was applied, with edge weights corresponding to cosine similarity scores and node colors representing BMN subtype. Node positions were computed with igraph’s default Fruchterman–Reingold force-directed layout, such that edges weighted by cosine similarity act as springs attracting similar neurons, while all nodes repel to avoid overlap. Consequently, neurons with higher similarity scores (i.e., similar postsynaptic connectivity profiles) are drawn close together, forming visually distinct clusters with varying overlap. This graph was used to visualize the spatial relationships among individual BMNs in similarity space.

Several versions of the analysis were performed, including matrices that incorporated synaptic weights and omitted cosine similarity thresholding. All approaches revealed consistent overall structure; however, the binarized matrix with similarity thresholding yielded the clearest organization and was selected for final visualization.

### Pre/post neurons as feedback loops

To investigate potential feedback loops between BMNs and their synaptic partners, we performed further analysis on partners that were both pre- and postsynaptic to BMNs (“pre/post” partners). As described above, synaptic edge lists were compiled from Codex data containing all BMN connections. These lists were then filtered to retain only connections between BMNs and the 39 non-BMN partners identified as both pre- and postsynaptic.

To more precisely define true pre/post partners and exclude those with highly asymmetric connectivity, we calculated the ratio of total presynaptic to postsynaptic connections for each partner. Only neurons with a ratio between 0.1 and 10 were retained, meaning that a partner could not be predominantly presynaptic or postsynaptic by more than one order of magnitude. This criterion allowed us to identify neurons with reasonably balanced bidirectional connectivity and improved the interpretability of the visualization. Five neurons were excluded based on this threshold, leaving 34 true pre/post partners in the final analysis (**Supplementary file 13**).

A network graph showing all connections between BMNs and the 34 true pre/post partners was generated using the igraph and visNetwork packages in R, as described above (**Figure 11 – figure supplement 1**). We further filtered the edge data to include only reciprocal connections between BMN subtypes and pre/post neurons. For each pair of BMN type and pre/post partner, both presynaptic and postsynaptic connections were required for the edge to be retained. All non-reciprocated edges (i.e., those where a partner is postsynaptic to one BMN type and presynaptic to another BMN type) were excluded. This filtered subset was visualized as a second directed network graph, highlighting bidirectional loops between specific BMN types and pre/post partners (**Figure 11A**).

### Identifying BMN postsynaptic hemilineages

Postsynaptic partners of BMNs were further annotated by developmental origin using hemilineage assignments from the classification.csv table in Codex (https://codex.flywire.ai/). This allowed each partner to be assigned to one of three categories: putative primary neurons, secondary hemilineage-derived neurons, or undefined/unassigned neurons. Predicted neurotransmitter identities for each partner were retrieved from the Codex neurons.csv table and merged with the hemilineage assignments (**Supplementary file 6**).

For each hemilineage, we counted the number of BMN-connected neurons per hemisphere and compared this to the total number of neurons in that hemilineage as reported by Schlegel *et al*. This allowed calculation of the fraction of each hemilineage that receives direct BMN input (**Figure 12A**). The complete list of connected hemilineages and their synapse counts is provided in **Supplementary File 14**.

Anatomical distributions of BMN-connected versus unconnected neurons within each hemilineage were visualized by mapping neuron meshes in FlyWire (**Figure 12B–T**, **Figure 12—figure supplement 1**). The overall composition of BMN postsynaptic partners across all partner categories (e.g., interneurons, descending neurons), subdivided by hemilineage and neurotransmitter, was summarized in **Figure 12—figure supplement 2**.

### Assessing relationship between BMNs and LB23 neurons

To explore the connectivity between BMNs and neurons of hemilinegage 23b (LB23), we constructed two network visualizations based on Codex-derived synapse data. Synaptic output data from BMNs (**Supplementary file 10**) were filtered to include only postsynaptic partners annotated as LB23 neurons (**Supplementary file 16**). The resulting matrix was used to generate a bipartite adjacency graph in which BMN types and LB23 neurons were represented as nodes, and the edges represented synaptic connectivity (**Figure 14B**).

The adjacency matrix was transposed and converted into a weighted directed graph using the igraph package in R, as described above. Node metadata were added from a manually curated annotation file (**Supplementary file 6**). Node shapes were assigned based on neuron categories (e.g. BMN, interneuron) and edge thickness was scaled to synapse count. The visIgraphLayout function was used to generate spatial node arrangements based on the force-directed layout algorithm by Fruchterman-Reingold, as described above.

In a second analysis, we focused on connections between BMNs and three individual antennal grooming neurons within the 23b lineage: aBN2_1, aBN2_2, and aBN2_3. Two rounds of filtering were performed on the BMN synaptic output matrix. First, the matrix was filtered to include only connections with the three aBN2 neurons, removing all other postsynaptic partners. Next, this reduced matrix was further filtered to retain only connections that met at least one of two criteria: (1) at least two BMNs from the same subtype were connected to the same aBN2 neuron, or (2) the connection contained more than five synapses. The resulting subset of connections was used to generate a simplified bipartite network graph, which was visualized using the visNetwork package with a hierarchical Sugiyama layout to highlight directional connectivity between BMN subtypes and aBN2 neurons (**Figure 14E**).

### Immunohistochemical analysis

We evaluated the expression pattern of the R40F04-GAL4 driver line by crossing it to 10XUAS-IVS-mCD8::GFP (RRID: BDSC_32194) (**Figure 13B**). The brains were dissected and stained as previously described (Hampel et al., 2015, 2011). For the staining we used anti-GFP and anti-nc82 with their respective secondary antibodies: rabbit anti-GFP (1:500, Thermo Fisher Scientific, Waltham, MA, #A11122), mouse mAb anti-nc82 (1:50,Developmental Studies Hybridoma Bank, University of Iowa), AlexaFluor-488 (1:500; goat anti-rabbit; Invitrogen, Carlsbad, CA), and AlexaFluor-568 (1:500; goat anti-mouse, goat anti-rat; Invitrogen).

For multicolor flipout (MCFO) experiments (**Figure 13D-F**), aBN2-spGAL4-1 (Hampel et al., 2015) was crossed to the MCFO-5 stock (RRID: BDSC_64089) (Nern et al., 2015). 1 to 3-day old fly brains were dissected and stained using anti-V5, -FLAG, and -HA antibodies. The following primary and secondary antibodies were used: rat anti-FLAG (Novus Biologicals, LLC, Littleton, CO, #NBP1-06712), rabbit anti-HA (Cell Signaling Technology, Danvers, MA, #3724S), mouse anti-V5 (Serotec, Kidlington, England #MCA1360), AlexaFluor-488 (1:500; goat anti-rabbit; Invitrogen), AlexaFluor-568 (1:500; goat anti-mouse; Invitrogen), AlexaFluor-633 (1:500; goat anti-rat; Invitrogen). We imaged individually labeled neurons from at least 10 brains.

Stained brains were imaged using a Zeiss LSM800 confocal microscope (Carl Zeiss, Oberkochen, Germany). Image preparation and adjustment of brightness and contrast were performed with Fiji software (http://fiji.sc/) (Schindelin et al., 2012). We compared the morphology of the imaged aBN2 neurons that were imaged via confocal microscopy with their corresponding EM reconstructed neurons using Fiji and Flywire, respectively.

## Acknowledgements

We thank Jonathan Blagburn for discussions and comments on this manuscript. We thank the Princeton FlyWire team and members of the Murthy and Seung labs for development and maintenance of FlyWire (supported by BRAIN Initiative grant MH117815 to Murthy and Seung), and the Princeton EM proofreading team for contributing neuronal edits. Research reported in this publication was supported by the National Institute Of Neurological Disorders And Stroke of the National Institutes of Health under Award Number R01NS121911. The content is solely the responsibility of the authors and does not necessarily represent the official views of the National Institutes of Health. This work was also supported by the Whitehall Foundation (2017-12-69), Puerto Rico Science, Technology & Research Trust (2020-00195), NIMHD MD007600 (RCMI), NIGMS-NIH RISE (R25GM061838), NIH NIGMS (P30-GM149367), and NSF (EES-1736019).

---

**Figure 1 – figure supplement 1.**
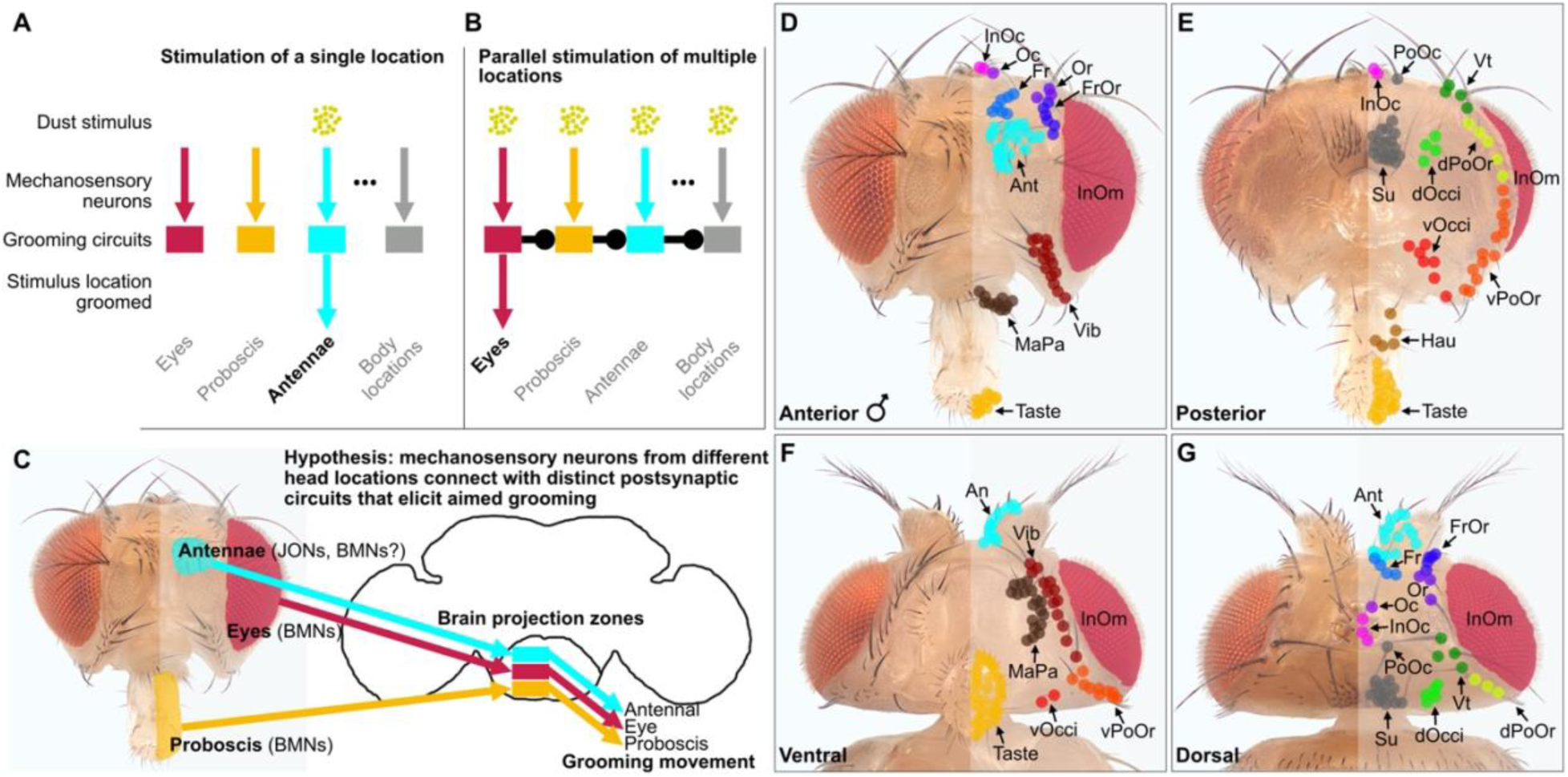
Proposed grooming circuit architecture suggests parallel mechanosensory pathways that are somatotopically organized. (**A,B**) This architecture facilitates aimed grooming of specific head and body locations (**A**) and controls hierarchical suppression during sequential grooming (**B**). (**A**) When a mechanical stimulus like dust touches a specific area (e.g., antennae), local mechanosensory neurons detect it. These neurons connect to excitatory postsynaptic circuits that elicit aimed grooming of those locations. This study emphasizes the connectome of bristle mechanosensory neurons (BMNs) responsible for grooming different head locations, such as the eyes (red), proboscis (orange), and antennae (aqua). The grooming pathway for body locations (gray) extends this parallel structure to any body part, like the abdomen or wings. (**B**) Dust covering multiple areas simultaneously causes competition among pathways that elicit exclusive grooming actions that are prioritized through hierarchical suppression (unidirectional connections between circuits shown). For clarity, only the nearest-neighbor connections are depicted. In this hypothesized structure, each circuit inhibits all lower-ranking circuits in the hierarchy. For instance, eye grooming occurs first as it inhibits all subsequent actions, like grooming the proboscis, antennae, and body areas. An alternative to the depicted unilateral inhibition is a sensory gain-mediated suppression mechanism (e.g. mechanosensory presynaptic inhibition). (**C**) Mechanosensory neurons from different head locations project to distinct, somatotopically organized zones in the ventral brain and elicit aimed grooming of those locations, including the antennae (via JONs [Johnston’s organ neurons; not analyzed in this study] and BMNs), eyes (BMNs), and proboscis (BMNs). The projections of mechanosensory neurons in the brain connect with postsynaptic circuits (indicated by boxes) that elicit grooming via descending pathways that activate movement pattern generators in the ventral nerve cord. (**D-G**) Bristles on the anterior (**D**), posterior (**E**), ventral (**F**), and dorsal (**G**) locations on the male head. The bristles on the right side are marked with color-coded dots for classification. Bristle full names are listed in the Figure 1 legend. Panels **A-C** were modified under the terms of the CCBY license from Figure 1 – figure supplement 1 of Eichler *et al*. (Eichler et al., 2024). Panels **D-G** were reproduced from Figure 1A-D of Eichler *et al*. (Eichler et al., 2024).

**Figure 1 – figure supplement 2.**
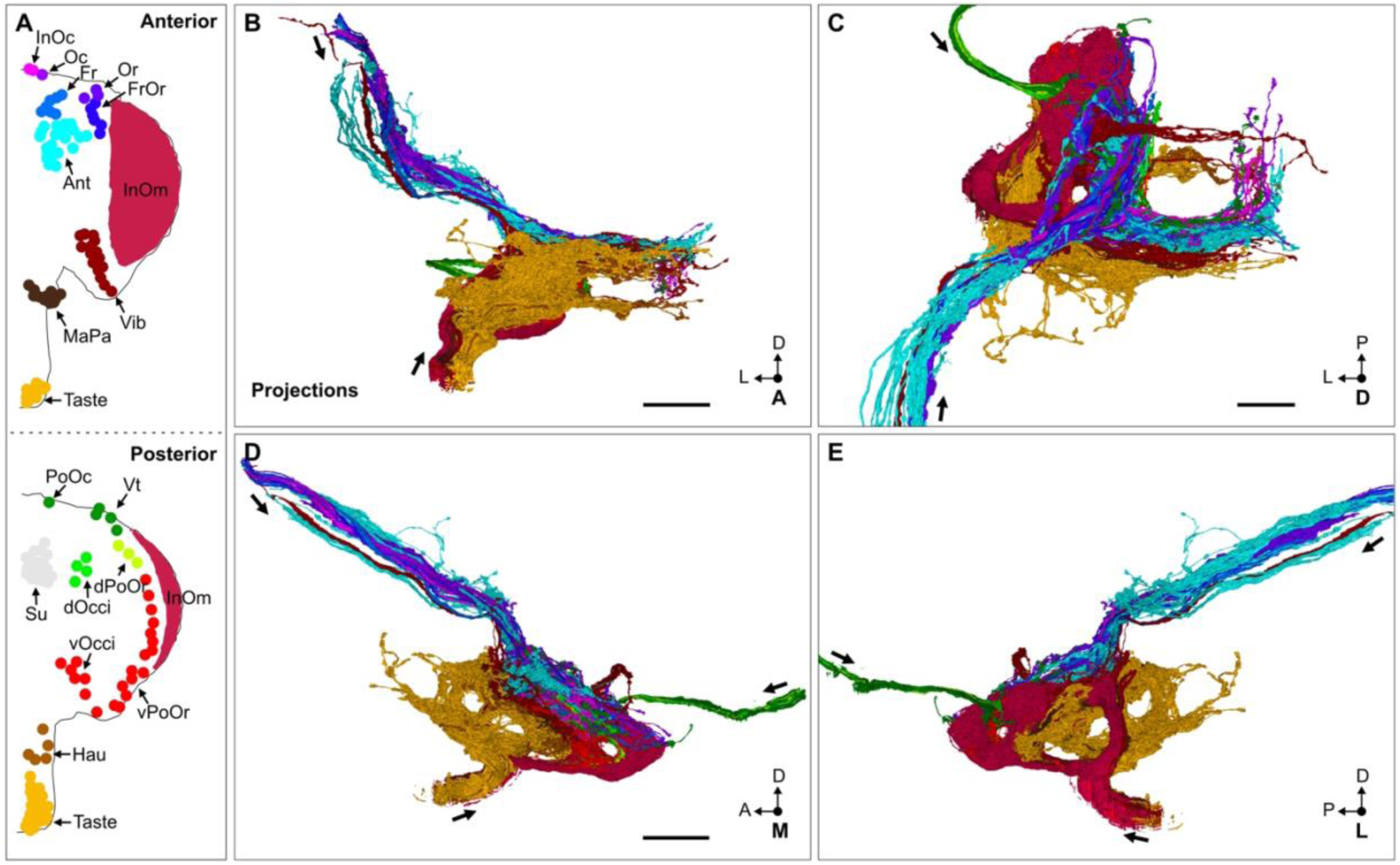
BMN type ventral brain projections. (**A**) Bristles of the anterior and posterior head. Bristle populations are marked with labeled and color-coded dots indicating their classification. Bristle population names are abbreviated (see Figure 1 legend for full names). (**B-E**) Reconstructed BMN projections colored by type according to the bristles that they innervate. Shown are the anterior (**B**), dorsal (**C**), medial (**D**), and lateral (**E**) views. Arrows indicate the projection directions for each incoming BMN nerve bundle. Scale bars, 20 μm. Panel **A** was modified under the terms of the CCBY license from Figure 6C,D of Eichler *et al*. (Eichler et al., 2024).

**Figure 1 – figure supplement 3.**
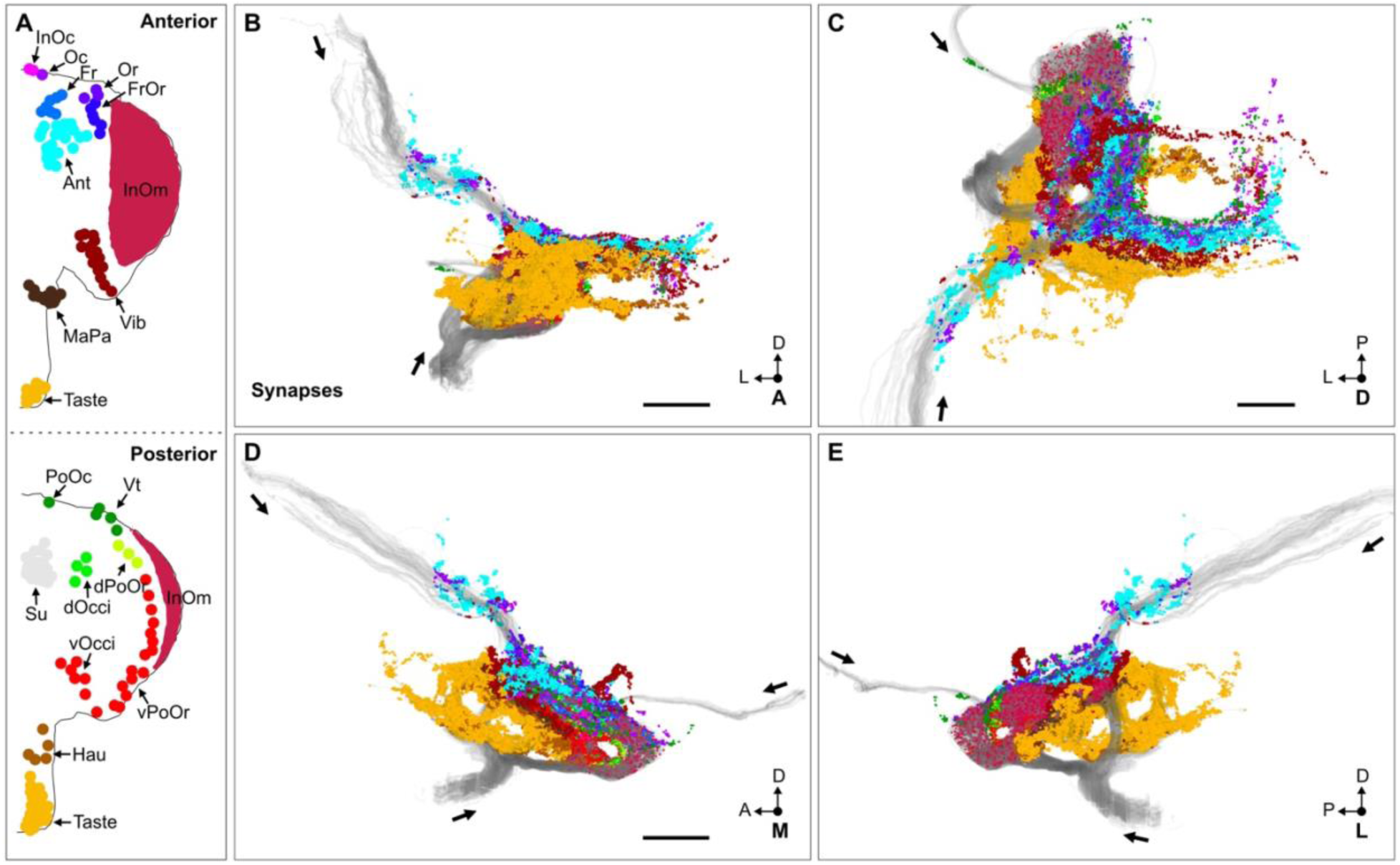
BMN type ventral brain synapses. (**A**) Bristles of the anterior and posterior head. Bristle populations are marked with labeled and color-coded dots indicating their classification. Bristle population names are abbreviated (see Figure 1 legend for full names). (**B-E**) Reconstructed BMN projections (gray) and their pre- and postsynaptic sites, colored by type according to the bristles that their corresponding BMNs innervate. Shown are the anterior (**B**), dorsal (**C**), medial (**D**), and lateral (**E**) views. These views highlight both overlapping projection zones, visible as intermingled synapses of different colors from neighboring BMN types, and segregated zones, where synapses from distinct BMN types remain spatially separated with minimal color mixing. Arrows indicate the projection directions for each incoming BMN nerve bundle. Scale bars, 20 μm. Panel **A** was modified under the terms of the CCBY license from Figure 6C,D of Eichler *et al*. (Eichler et al., 2024).

**Figure 2 – figure supplement 1.**
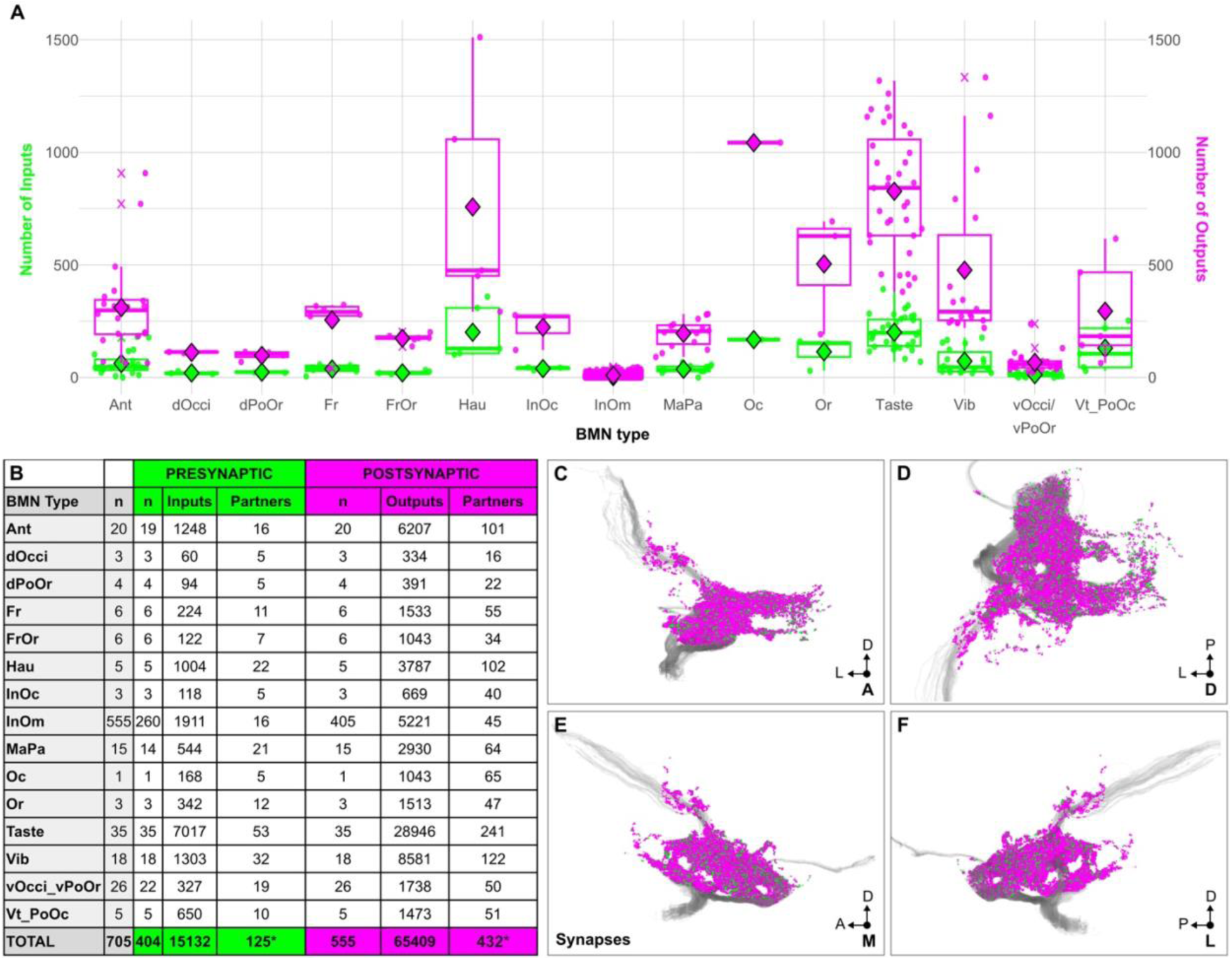
Distribution of BMN synapses. **(A)** Box plots of BMN pre- and post synapses by type. Each point (dot) indicates the number of synaptic inputs- (green) or outputs (magenta) for an individual BMN. Points are horizontally dispersed for easier visualization. Box indicates the interquartile range (IQR) for the particular BMN type, horizontal bar in box indicates the median, diamond indicates the mean, and vertical lines extend to furthest points within 1.5 times the IQR. “X” indicates outliers. (**B**) Table indicating cumulative number of inputs, outputs, and partners for each BMN type. n in gray indicates the number of left side BMNs that were reconstructed (Eichler et al., 2024). n in green and magenta indicate the number of BMNs that were connected with at least 5 synapses to any given pre- or postsynaptic neuron, respectively. The number of inputs or outputs refers to synapses, while the number of partners refers to pre- or postsynaptic neurons. *The total number of pre- or postsynaptic partners is lower than the total of connected partners because some partners are connected with multiple BMN types. Underlying postsynaptic and presynaptic data is in **Supplementary file 2** and **Supplementary file 3**, respectively. (**C-F**) BMN synapse locations, colored by BMN inputs (green) and outputs (magenta); shown in anterior (**C**), dorsal (**D**), medial (**E**), and lateral (**F**) views. Reconstructed BMNs are shown in gray.

**Figure 4 – figure supplement 1.**
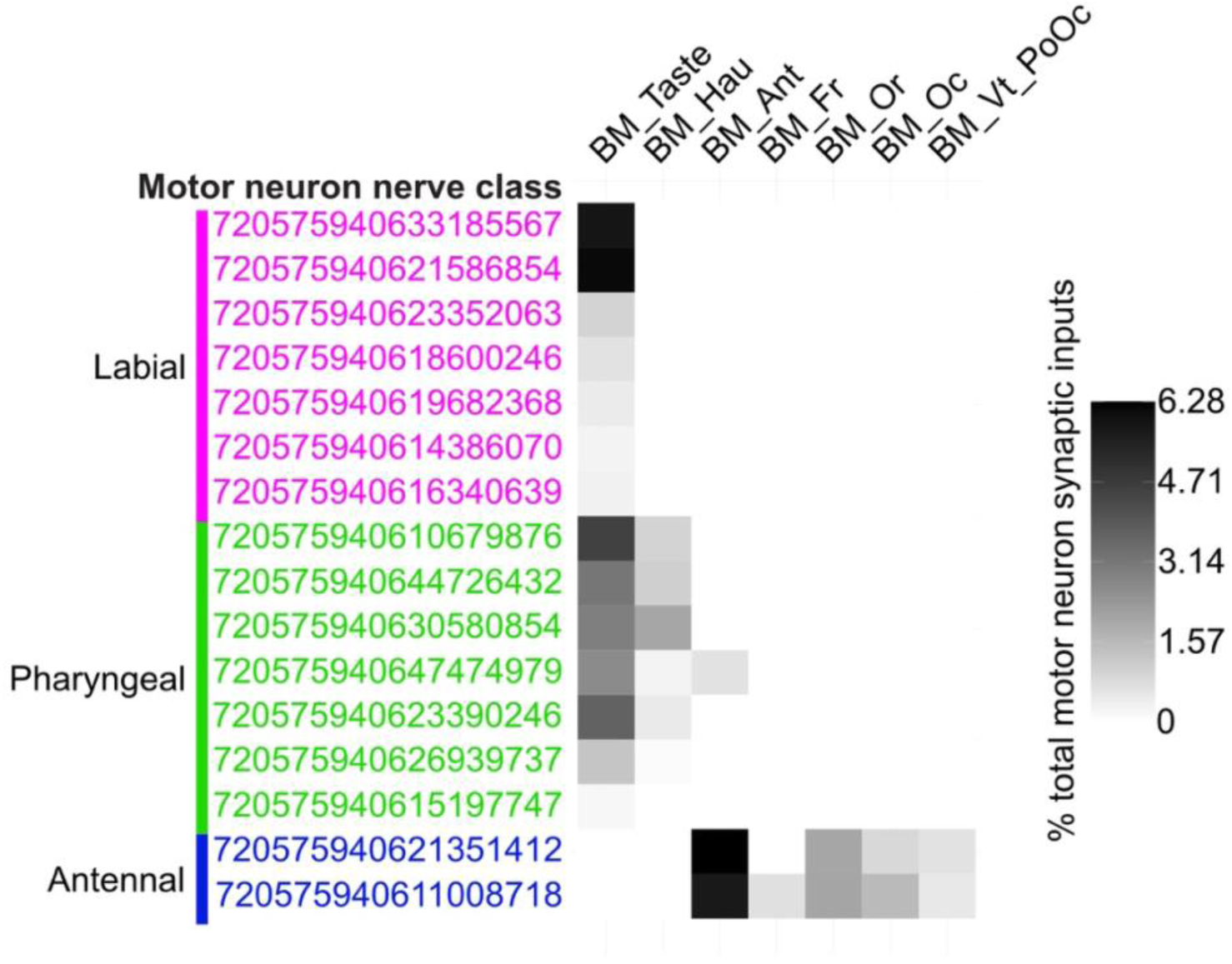
Percent of the total motor neuron synaptic inputs from each BMN type. Heatmap indicating the percent of the total synaptic inputs from any neuron that are from each BMN type. Grayscale shading maximum indicates that BM-Ant neurons occupy 6.28% of the total presynaptic sites onto antennal nerve class neuron 720575940621351412. Underlying data are in **Supplementary file 7**. Note: to observe the extent of BMN synaptic inputs into the motor neurons, the analysis shown here includes reconstructed BMNs from the left and right sides of the head. Left side BMNs are shown in Figure 3C. Right side BMN types include BM-Taste, -Hau, -Ant, -Fr, -Or, -Oc, and -Vt/PoOc neurons, of which the BM-Fr neurons are the only type that are uniquely connected from the right side but not the left.

**Figure 11 – figure supplement 1.**
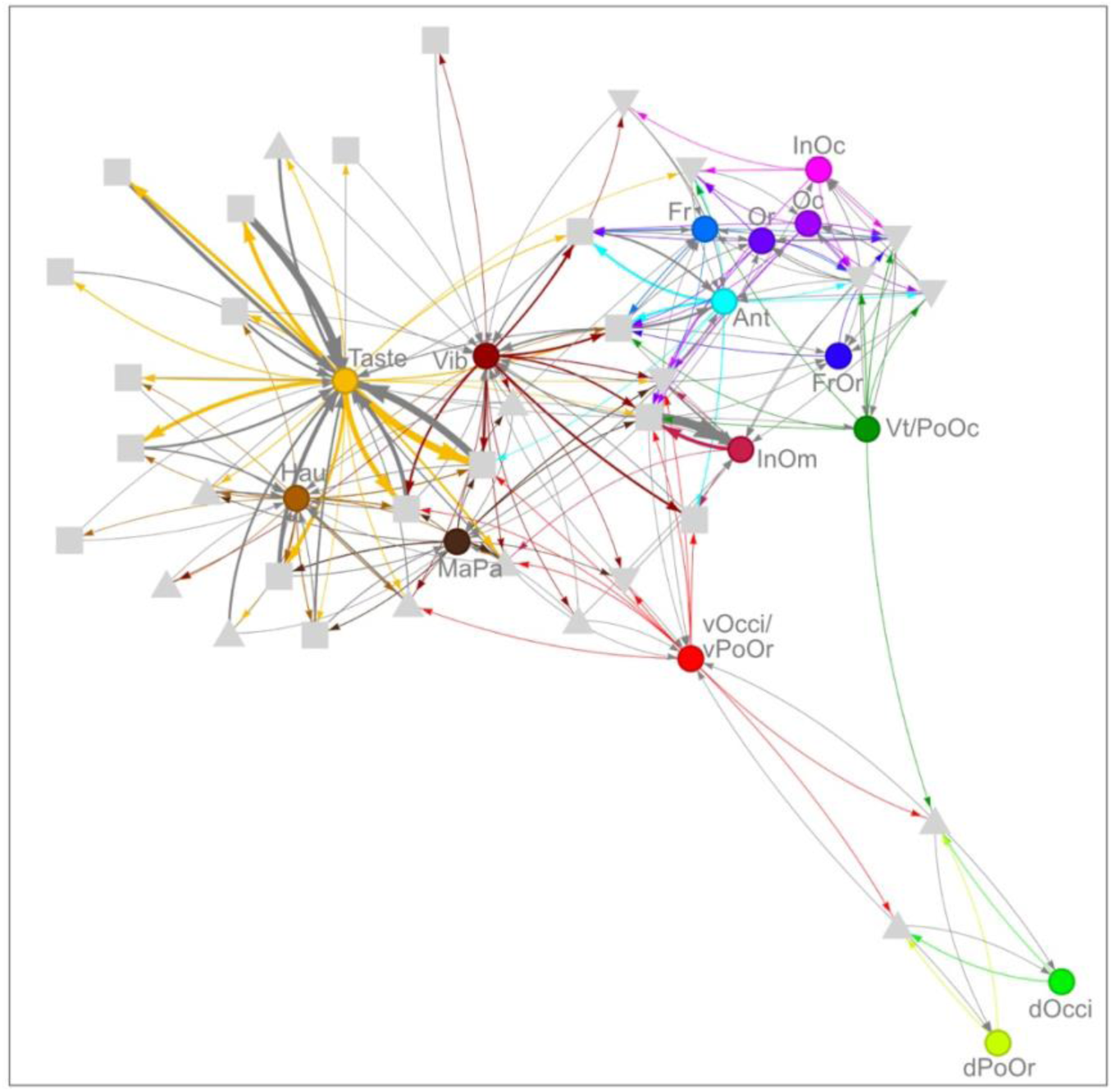
BMN connectome with pre/post neurons shows somatotopic organization. (**A**) Network diagram of 34 pre/post neurons whose pre:post ratio is within 0.1-10 (gray) with different BMN types projecting into the left brain hemisphere (colored dots). Squares correspond to individual interneurons while down and up triangles indicate descending and ascending neurons, respectively. Edges are in colors corresponding to the BMN type outputs, while gray edges correspond to BMN GABAergic inputs. Node positions were computed using a Fruchterman–Reingold force-directed layout: each directed edge acts like a spring whose strength is proportional to the edge weight (synapse count), drawing highly connected nodes together, while all nodes repel one another to prevent overlap. Consequently, neurons with many or strong synaptic connections cluster closely in the 2D embedding, whereas weakly or unconnected neurons are pushed to the periphery. Underlying data are in **Supplementary file 13** which includes a complete account of the pre/post neuron connections with the BMN types.

**Figure 12 – figure supplement 1.**
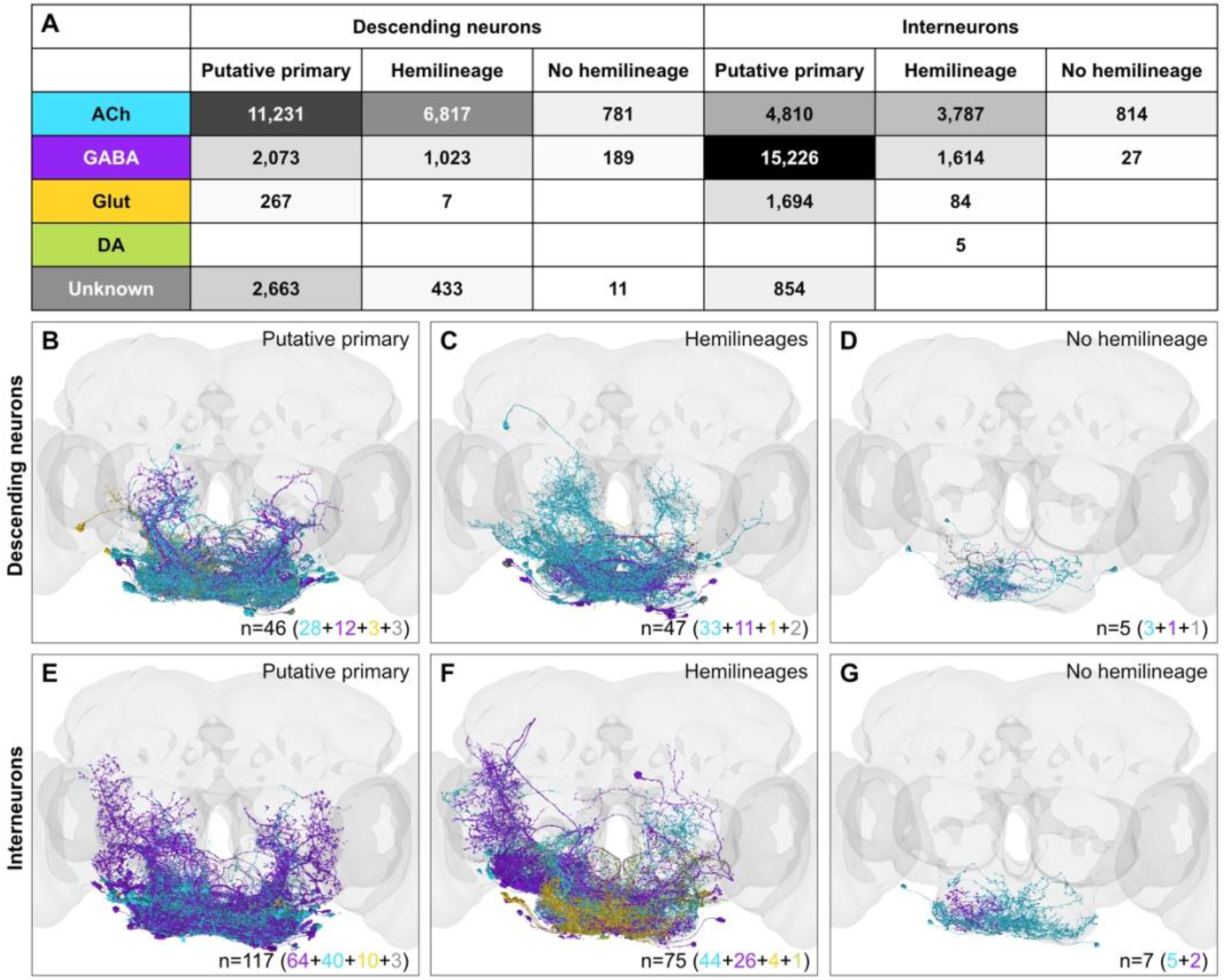
BMN postsynaptic partner developmental origins. (**A**) BMN synapse numbers onto postsynaptic partners that use different neurotransmitters. Table indicates the number of BMN synapses onto postsynaptic descending neurons and interneurons. These neurons are subdivided based on developmental birth timing (Schlegel et al., 2024), including putative primary neurons of embryonic origin, part of a postembryonic hemilineage, or those with no hemilineage assigned. Partners are predicted to use acetylcholine (ACh), GABA, glutamate (Glut), or dopamine (DA). Unknown indicates synapses whose neurotransmitter identities could not be determined by the prediction algorithm (Eckstein et al., 2024). Grid grayscale shades indicate synapse numbers, with black indicating 15,226 synapses. Neurotransmitter color codes are used in **B-G**. (**B-G**) Anterior views of BMN postsynaptic descending neurons (**B-D**) and interneurons (**E-G**) belonging to putative primary neurons (**B,E**), hemilineage neurons (**C,F**), and neurons with no hemilineage assigned (**D,G**). Colors indicate the predicted neurotransmitters for each partner, including GABA (purple), ACh (teal), Glu (mustard), and DA (green). n indicates the total number of neurons in each subdivision. Colors in parentheses represent predicted neurotransmitter identities using the color code from panel (**A**). Underlying data are in **Supplementary file 14**.

**Figure 12 – figure supplement 2.**
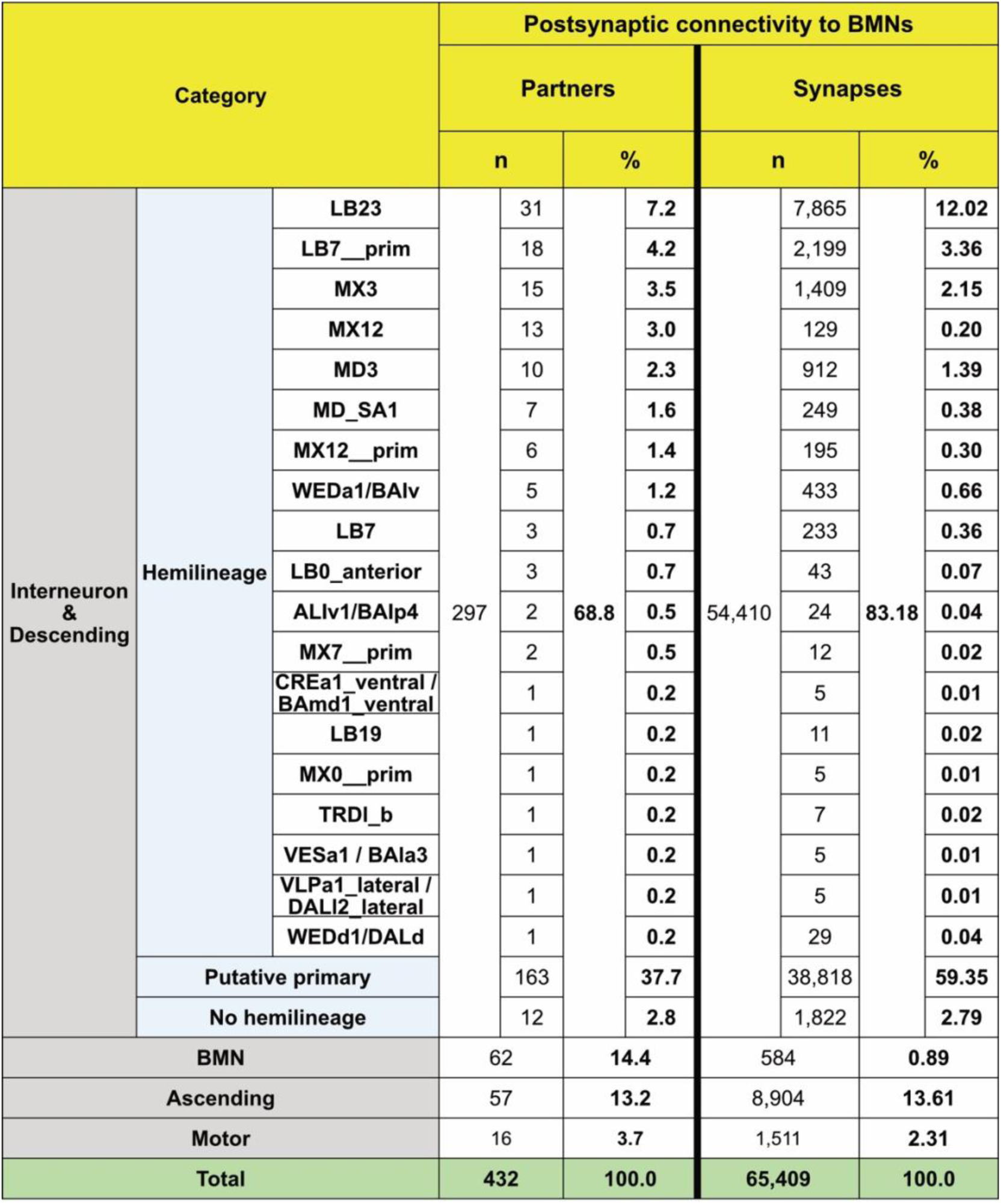
BMN postsynaptic partners. Neurons in each postsynaptic partner category and their connectivity to head BMNs. Left columns: different categories of partners including interneurons, descending neurons, BMNs, ascending neurons, and motor neurons. Interneurons and descending neurons are subcategorized as putative primary, assigned to a hemilineage, or not assigned to a hemilineage. Those assigned to a hemilineage are further subcategorized by the individual hemilineage they belong to. Centercolumns: n indicates the number of neurons in each category found to be postsynaptic to BMNs, expressed also as the percentage (%) of total BMN partners accounted for by each category (i.e., [n / 432]*100). Right columns: n indicates the total number of BMN output synapses corresponding to each category, expressed also as the percentage (%) of total BMN outputs accounted for by each category (i.e., [n / 65409]*100) . Underlying data are in **Supplementary file 14**.

**Supplementary file 1. Table of EM-reconstructed head bristle mechanosensory neurons (BMNs).** Table with individual BMNs as separate rows, with columns for BMN type, and number of inputs and outputs found using CODEX data.

**Supplementary file 2. Edge list of BMNs with postsynaptic partners.** Edge list with each row reflecting a connection between an individual BMN and a postsynaptic partner, and the number of synapses between them.

**Supplementary file 3. Edge list of BMNs with presynaptic partners.** Edge list with each row reflecting a connection between a presynaptic partner and an individual BMN, and the number of synapses between them.

**Supplementary file 4. Raw data: combined input/output adjacency matrix.** BMNs are grouped by type, and partners double-grouped by category (ascending, descending, etc.) and relationship (presynaptic, postsynaptic, or pre/post).

**Supplementary file 5. Normalized data: combined input/output adjacency matrix.** BMNs grouped by type, and partners double-grouped by category (ascending, descending, etc.) and relationship (presynaptic, postsynaptic, or pre/post). Data normalized by dividing each value by total outputs for that BMN type. This data was used to make the sankey diagram in **Figure 3A**. Normalization enables uniform length of rectangles representing BMNs in panel **A**.

**Supplementary file 6. Summary of BMN partners.** Table containing a row for each BMN partner containing its Flywire ID, category, neurotransmitter information, and number of times pre- and/or postsynaptic to BMNs.

**Supplementary file 7. Connections of different BMN types with motor neurons.** Numbers of BMNs of each type that are connected with each motor neuron, used to generate **Figure 4C**. Also, calculated percentages of the total motor neuron synaptic inputs from each BMN type, used to generate **Figure 4 – figure supplement 1**.

**Supplementary file 8. BMN partner synapses in different brain neuropils.** Table with every pre, post, and pre/post partner, along with number of all input/output synapses in each neuropil of the brain.

**Supplementary file 9. BMN partner synapses in different brain neuropils (top 31 neuropils) and their normalized values.** BMN partner pre- and postsynapse counts for the 31 neuropils with the highest synaptic counts (total synapse count > 100). Shown are counts for pre- and postsynaptic, and pre/post partners. Top 31 neuropils, separated into columns for partners, further divided into input or output. Each column has a corresponding column containing the normalized values used for visualization.

**Supplementary file 10. BMN connectome edge list.** Edge list containing all connections from presynaptic partners to BMNs, connections from BMNs to postsynaptic partners, and their respective weights.

**Supplementary file 11. Individual BMN output edge list.** Edge list containing all connections between individual BMNs and their postsynaptic targets with their respective weights.

**Supplementary file 12. Matrix of individual BMNs and their pairwise cosine similarity scores.** A 555 x 555 matrix of individual BMNs and their calculated pairwise cosine similarity scores.

**Supplementary file 13. Edge list of BMN connections with pre/post neurons.** Edge list containing all connections from pre/post neurons to BMNs, connections from BMNs to pre/post partners, and their respective weights. Provides additional information used for filtering based on pre:post ratio and reciprocal connectivity.

**Supplementary file 14. Developmental origin of BMN postsynaptic partners including hemilineage information.** Table of BMN postsynaptic and pre/post partners along with the hemilineage information. Provides additional information regarding total number of neurons within each hemilineage population and the proportion of these neurons that are postsynaptic targets of BMNs.

**Supplementary file 15. Counts of LB23 neurons in R40F04-GAL4.** Counts from the left and right brain hemispheres in nine individual brains of R40F04-GAL4 expressing mCD8::GFP.

**Supplementary file 16. FlyWire.AI IDs for different LB23 neurons.**

## Notes

### Competing Interest Statement

The authors have declared no competing interest.

### Summary of Updates

This revision addresses reviewer comments at the journal eLife. This draft will be resubmitted to the journal.

